# Multiunit frontal eye field activity codes the visuomotor transformation, but not gaze prediction or target memory, in a delayed saccade task

**DOI:** 10.1101/2023.10.08.560888

**Authors:** Serah Seo, Vishal Bharmauria, Adrian Schütz, Xiaogang Yan, Hongying Wang, J. Douglas Crawford

## Abstract

Single-unit (SU) activity − action potentials isolated from one neuron — has traditionally been employed to relate neuronal activity to behavior. However, recent investigations have shown that multi-unit (MU) activity − ensemble neural activity recorded within the vicinity of one microelectrode − may also contain accurate estimations of task-related neural population dynamics. Here, using a well-established model-fitting approach, we compared the spatial codes of SU response fields with corresponding MU response fields recorded from the frontal eye fields (FEF) in head-unrestrained monkeys (*Macaca mulatta*) during a memory-guided saccade task. We focused on characterizing the visuomotor transformation from Target-in-Eye coordinates to future Gaze-in-Eye coordinates (Sajad et al., 2015). Most SU *visual* response fields coded targets (with some predicting Gaze), whereas the MU population only coded targets. Most SU *motor* responses coded Gaze, but many still retained a target code. In contrast, MU motor activity predominantly coded Gaze with very little target coding. Finally, both SU and MU populations showed a progressive transition through intermediate ‘Target-to-Gaze’ codes during the delay period, but the MU activity showed a ‘smoother’ transition. These results confirm the theoretical and practical potential of MU activity recordings as a biomarker for fundamental sensorimotor transformations (e.g., Target-to-Gaze coding in the oculomotor system), while also highlighting the importance of SU activity for coding more cognitive (e.g., predictive / memory) aspects of sensorimotor behavior.

**SIGNIFICANCE STATEMENT:** Multi-unit recordings (undifferentiated signals from several neurons) are relatively easy to record and provide a simplified estimate of neural dynamics, but it is not clear which single-unit signals are retained, amplified, or lost. Here, we compared single-/multi-unit activity from a well-defined structure (the frontal eye fields) and behavior (memory-delay saccade task), tracking their spatial codes through time. The progressive transformation from target to gaze coding observed in single-unit activity was retained in multi-unit activity, but gaze prediction (in the visual response) and target memory (in the motor response) were lost. This suggests that multi-unit activity provides an excellent biomarker for healthy sensorimotor transformations, at the cost of missing more subtle cognitive signals.

## INTRODUCTION

In systems neuroscience, it is common to estimate neuronal population dynamics by relating recordings of neuronal activity to some concurrent task or behavior. Traditionally, neuroscientists have isolated single-unit (SU) action potentials of individual neurons from raw recorded background noise, based upon its unique spike shape; a process called spike-sorting. Recently, however, there has been increased interest in the use of multi-unit (MU) activity (i.e., unsorted signals recorded from the tip of an electrode, which may arise from several nearby neurons (Ayar et al., 2023; Buzsáki, 2004; Panzeri et al., 2015; Quian Quiroga and Panzeri, 2009; Rossi-Pool and Romo, 2019). It has been suggested that MU activity recordings have practical and theoretical implications for experimental and applied electrophysiology (Ahmadi et al., 2021; Choi et al., 2010; Fraser et al., 2009; Leuthardt et al., 2021; Mattia et al., 2010; Sharma et al., 2015; Smith et al., 2013; Trautmann et al., 2019; Zhang and Constandinou, 2021). However, it remains unclear what is highlighted, and what is obliterated, when one combines the activity of several neurons into one conglomerate signal. Our aim was to test this question in a well-defined sensorimotor system (the frontal eye fields, FEF), task (memory-delay saccades), and analytic technique, as described in detail below.

Although SU recordings remain important for many applications, they pose both practical and theoretical challenges. The detection and extraction of SU amongst a sea of extracellular signals is often hindered by variability in spike shape / amplitude, and extraneous background noise (Lewicki, 1994; Mtetwa and Smith, 2006). Isolation of single action potentials is further confounded by overlapping spikes, making it hard to identify and classify spikes (Mtetwa and Smith, 2006; Quiroga et al., 2004; Rossi-Pool and Romo, 2019). These are time-consuming procedures that normally can only be done after recordings are complete. Further, neurons do not work in isolation, but as ensembles forming functional networks (Buzsáki, 2010; Edelman, 1987; Hebb, 1949; Levy et al., 2020; Miller et al., 2014; Molotchnikoff et al., 2019; Singer, 2013).

MU activity analysis sidesteps these challenges by avoiding the need to identify unique spike waveforms. The process of extracting / sorting MU activity is more reproducible since it does not eliminate data or depend on specific sorting techniques / parameters. Some studies found that MU activity provided a more stable signal with higher spatial resolution than SU activity (Ayar et al., 2023; Drebitz et al., 2019; Lewicki, 1998; Sharma et al., 2015; Xing et al., 2009). Further, MU activity profiles are similar to local field potential (LFP) signals but tend to show less correlation between nearby neurons (Ahmadi et al., 2021; Burns et al., 2010; Drebitz et al., 2019; Telenczuk and Destexhe, 2020). Finally, MU analysis is capable of decoding the task and behavior by reducing the dimensionality of neural states (Ayar et al., 2023; Quian Quiroga and Panzeri, 2009; Rossi-Pool and Romo, 2019; Smith et al., 2013; Trautmann et al., 2019). Therefore, MU activity analysis has become popular for applications such as brain-machine interfaces, recording paralytic motor activity (Gilja et al., 2015; Pandarinath et al., 2017) and other clinical / preclinical trials (Christie et al., 2015; Fraser et al., 2009; Kao et al., 2017; Todorova et al., 2014). Some research suggests that MU activity sometimes contains more accurate estimates of neural population dynamics (Sharma et al., 2015; Trautmann et al., 2019). However, it remains unclear which signals are retained, and which are lost, when one switches from SU to MU analysis.

The gaze control system provides an ideal model system to compare the SU *vs.* MU codes, because spatiotemporal codes have already been described at the SU level. Neurons in higher-level gaze structures [superior colliculus (SC), lateral intraparietal cortex (LIP), frontal eye fields (FEF) and supplementary eye fields (SEF)] show separate visual and motor responses when a memory delay separates visual target presentation and saccade initiation (Gnadt et al., 1991; Heusser et al., 2022; Schall, 2015; Umeno and Goldberg, 2001; White et al., 1994). Further, these responses are spatially organized into visual and motor response fields (Andersen et al., 1992; Hafed et al., 2023; Schall, 2015; Schlag and Schlag-Rey, 1987; Sommer and Wurtz, 2001). Finally, although results vary with task, higher-level visuomotor structures primarily show eye-centered response fields (Andersen et al., 1985; Caruso et al., 2018a, 2018b; Constantin et al., 2007; Keith et al., 2009; Mullette-Gillman et al., 2005; Park et al., 2006; Snyder, 2005). For example, the FEF and SEF, located in dorsolateral and medial frontal cortex, show predominantly eye-centered sensory / motor response fields for visual targets / saccades respectively (Peck et al., 1995; Caruso et al., 2021).

We have used a model-fitting approach (fitting various spatial models against response field data) to track sensorimotor transformations in the gaze system through time (Keith et al., 2009; Sadeh et al., 2020, 2015; Sajad et al., 2016, 2015). In contrast to decoding techniques, which allow one to extract information from noisy neural populations (Fraser et al., 2009; Meyers, 2013; Bremmer et al., 2016; Brandman et al., 2017; Dong et al., 2023), this technique tests the original spatial code used by individual neurons or populations (Ahmadi et al., 2021; Burns et al., 2010; Drebitz et al., 2019; Keith et al., 2009). Initial investigations in SC and FEF confirmed a distributed, eye-centered transformation from target (Te) coding in visual response fields to future gaze (Ge) coding in the saccade motor response. When a spatial ‘T-G’ continuum between Te and Ge was used to track these codes through time, they showed a progressive transition from target to gaze coding at the population level (Sadeh et al., 2018, 2015; Sajad et al., 2016, 2015), even without a memory delay (Sadeh et al., 2020). But not all cells showed this transition: some visual response predicted Ge and some motor responses retained Te. Recently, this technique was applied to FEF / SEF response fields in the presence of background landmarks (**Fig. 1A).** Although background stimuli produced various modulations (Schütz et al., 2023), the data still showed a strong T-G transformation (Bharmauria et al., 2021, 2020). Therefore, we deemed this might provide an ideal dataset to test the relative robustness of sensorimotor codes derived from SU versus MU activity.

**Figure 1:**
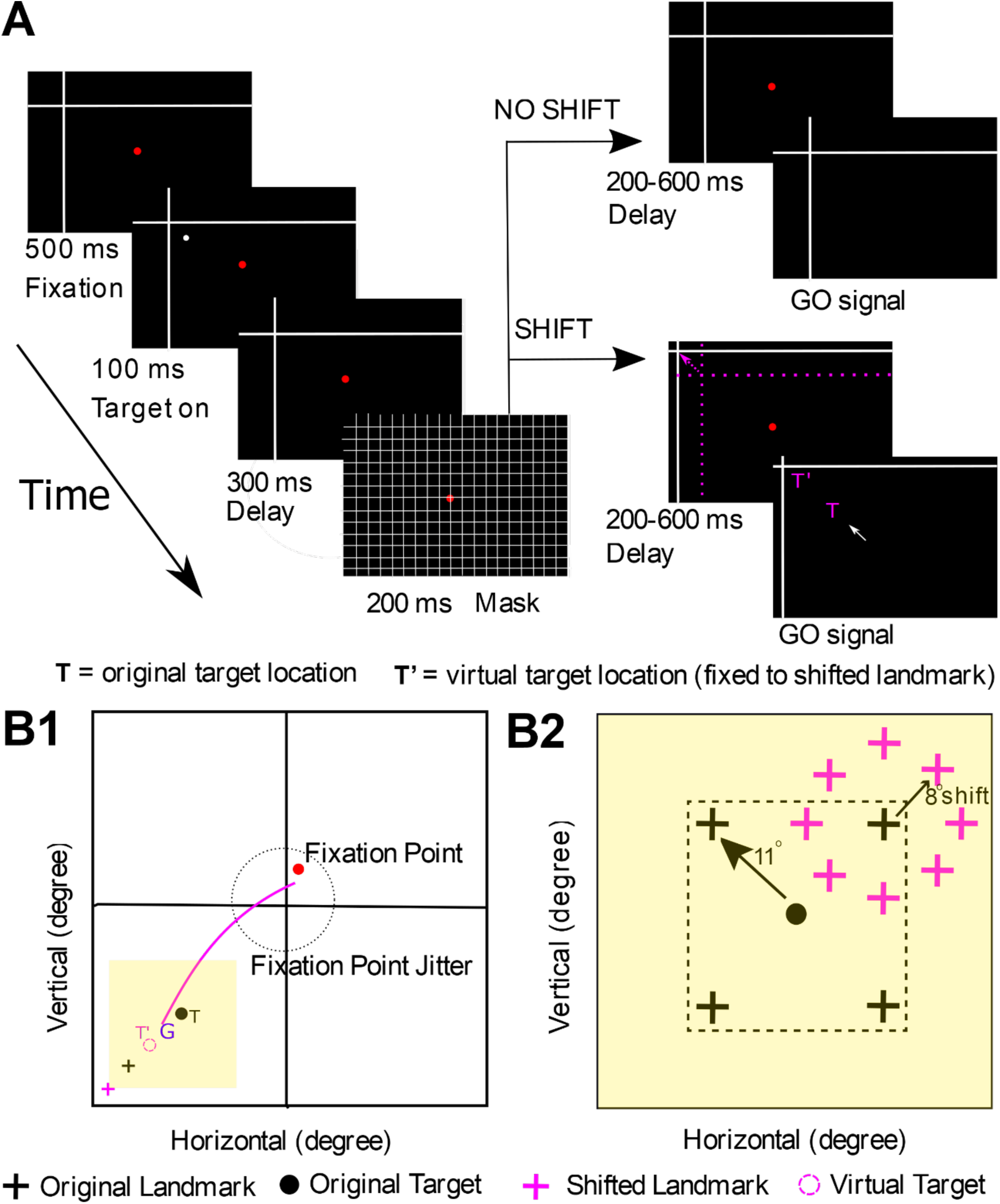
Representation of the experimental task and behavior. **A**, Cue-conflict paradigm, and its time course. The monkey began the task by fixating on a red dot for 500 ms while a landmark (white, intersecting lines) was present on the screen. Next, a target (white dot) was flashed for 100 ms, followed by a 300 ms memory delay where the target disappeared from the screen. After a 200 ms grid-like mask and another memory delay (200-600 ms), the monkey was cued by the disappearance of the fixation dot (i.e., go signal) to saccade head-unrestrained toward the remembered target location either when the landmark was shifted (represented by the broken pink arrow) or when it was not shifted. The monkey was rewarded when they placed their gaze (G) within a radius of 8-12° around the original target [at T (original target), T’ (virtually shifted target fixed to shifted landmark) or between T and T’]. The reward window was centered on T so that behavior was not biased. Note that the actual landmark shift was 8°, but it has been exaggerated in the schematic figure for clarity. The pink-colored objects are only used in the schematic figure for purposes of representation. **B1**, Representation of a gaze shift (pink curve) from the initial fixation point (red dot) toward the virtual target (dotted pink circle) fixed to shifted landmark (pink cross). G represents the final gaze location, and the yellow square represents the area in B2. The dotted circle stands for the jitter of the initial fixation points on the screen from one trial to another. **B2**, Representation of all (four) possible landmark (black cross) locations and shifted landmark (pink cross, relative to the original landmark location) locations for an example target (black dot). The landmark was presented 11° around the original target in one of four oblique directions. The shifted landmark was presented 8° around the original landmark location in one of eight radial directions.

Here, we analyzed the spatial codes of SU and MU activity derived from the same FEF recording sessions (Bharmauria et al., 2020). After isolating SU and MU activity (Drebitz et al., 2019; Quiroga et al., 2004), we applied our model-fitting approach to test their visual and motor response fields. After confirming that SU / MU visual and motor population codes were best characterized overall by the Te and Ge models respectively, we used the ‘T-G’ spatial continuum between these models to characterize, quantify and compare the spatiotemporal progression of SU *vs.* MU activity coding schemes (DeSouza et al., 2011; Keith et al., 2009; Sadeh et al., 2015; Sajad et al., 2015). We found that the MU and SU response fields show similar spatial codes and spatiotemporal transformations, but MU showed a ‘smoother’ sensorimotor transition and lacked certain nuances (i.e., gaze prediction in visual responses and residual target signals in motor responses), resulting in a ‘cleaner’ sensorimotor transition. We conclude that MU activity could provide a practical, quick, and reliable biomarker for basic sensorimotor transformations but may lack certain nuances important for the more cognitive aspects of sensorimotor behavior.

## MATERIALS AND METHODS

Most experimental and analytic details of this study were published previously in analysis of SU response fields in the FEF (Bharmauria et al., 2020). These are summarized here along with specific details related to the MU activity response field analysis.

### Surgical Procedures

All experimental protocols were approved by the York University Animal Care Committee and were in alignment with the guidelines of Canadian Council on Animal Care on the use of laboratory animals. Neural data used in this study were gathered from two female *Macaca mulatta* monkeys (Monkey V and Monkey L). Two 3D search coils, each with a diameter of 5 mm, were implanted in the sclera of the left eye of the respective animal. The recording chambers in the FEF of both animals were implanted and centered at 25 mm anterior and 19 mm lateral. Surgeries were performed as previously done (Crawford et al., 1999; Klier et al., 2003). There was a craniotomy (19 mm diameter) beneath each chamber that allowed access to the right FEF. During the experiment, animals were placed in custom-designed chairs which allowed free head movements. To prevent animals from rotating in the chair, they wore a vest fixed to the chair. Furthermore, animals were placed in the setup with three orthogonal magnetic fields with two orthogonal cells mounted on their heads (Crawford et al., 1999). These fields induced a current in each coil. The amount of current induced by one of the fields is proportional to the area of the coil parallel to this field. This allowed us to derive the orientation of each coil in relation to the magnetic fields, as well as other variables including eye orientations, head velocities and eye and head accelerations (Crawford et al., 1999)

### Behavioral Paradigm

We chose the following paradigm for the current study because it provides a visually rich, noisy dataset for extraction of spatial codes and has already been applied to an existing dataset of FEF recordings. We employed head-unrestrained gaze shifts (i.e., including coordinated eye saccades and head motion) to separate signals organized relative to eye, head, or space coordinates. Visual stimuli were presented using laser projections on a flat screen that was placed 80 cm away from the animal **(Fig. 1A).** The monkeys performed a memory-guided gaze task in which animals were trained to remember a target relative to a visual allocentric landmark (displayed using two intersecting lines). This led to a temporal delay between the target presentation and eye movement initiation. This allowed us to independently analyze visual (aligned to target) and eye-movement-related (saccade onset) responses in the FEF. The experiments were conducted in a dark room to prevent any extraneous allocentric cues. Each trial began with the animal fixating on a red dot in the center of the screen for 500 ms in the presence of a landmark. A brief flash of the visual target (T, white dot) followed for 100 ms, then a 300 ms delay, followed by a grid-like mask for 200 ms (this concealed past visual traces, and current and future landmarks) and a second memory delay (200-600 ms). The disappearance of the red fixation dot signaled the animal to perform a head-unrestrained saccade (indicated by the solid white arrow) toward the memorized location of the target either in the presence of a shifted landmark (broken pink arrow, 90 % of trials shifted in one of the eight radial directions by 8°) or a non-shifted landmark (10 %, zero-shift condition, i.e., landmark position did not change as before mask). The saccade targets were flashed one-by-one in a random manner throughout a neuron’s response field. The details of the task are outlined in **Figure B1-B2**. In the shift condition, where the shifted target (T’) was fixed to the landmark, the animal was rewarded with a water-drop when its gaze (G) landed within 8-12° radius of the original target (animals were rewarded if they looked at T, T’, or anywhere in between). This large reward window is what allowed variability in the memory-guided gaze shifts and error accumulation (Gnadt et al., 1991; Sajad et al., 2015; White et al., 1994), and thus formed the logical base of our analysis (see below).

The response fields were tested roughly across horizontal and vertical dimensions (rectangular range of 30-80°). For both animals, there was a high correlation between the direction of the target and the gaze in both dimensions. This implies that that their gaze was aimed toward the vicinity of the T.

### Electrophysiological and Behavioral Recordings

We recorded acutely from the FEF, a gaze area associated with sensorimotor transformations (Caruso et al., 2018b; Siegel et al., 2015). Extracellular activity was recorded by lowering tungsten electrodes (0.2–2.0 mΩ impedance, FHC Inc.) into the FEF [using Narishige (MO-90) hydraulic micromanipulator] using the Plexon MAP System. The recorded activity was then digitized, amplified, filtered, and saved for offline spike sorting (using template matching). The recorded sites (in head-restrained conditions) were further confirmed using a low-threshold electrical microstimulation (50 μA) as used previously (Bruce et al., 1985). A superimposed pictorial of the recorded sites from both animals is presented in **Figure 2A-B** (Monkey L in Blue and Monkey V in red). **Figure 2C-D** displays the overlapped and averaged spike density plots for top 10 % (for each neuron and MU site) trials for the SU and MU. An average of 331 ± 156 (mean ± SD) trials/neuron were recorded. Mostly, the monkeys were free to scan the environment (head-unrestrained) while neurons were searched for.

**Figure 2:**
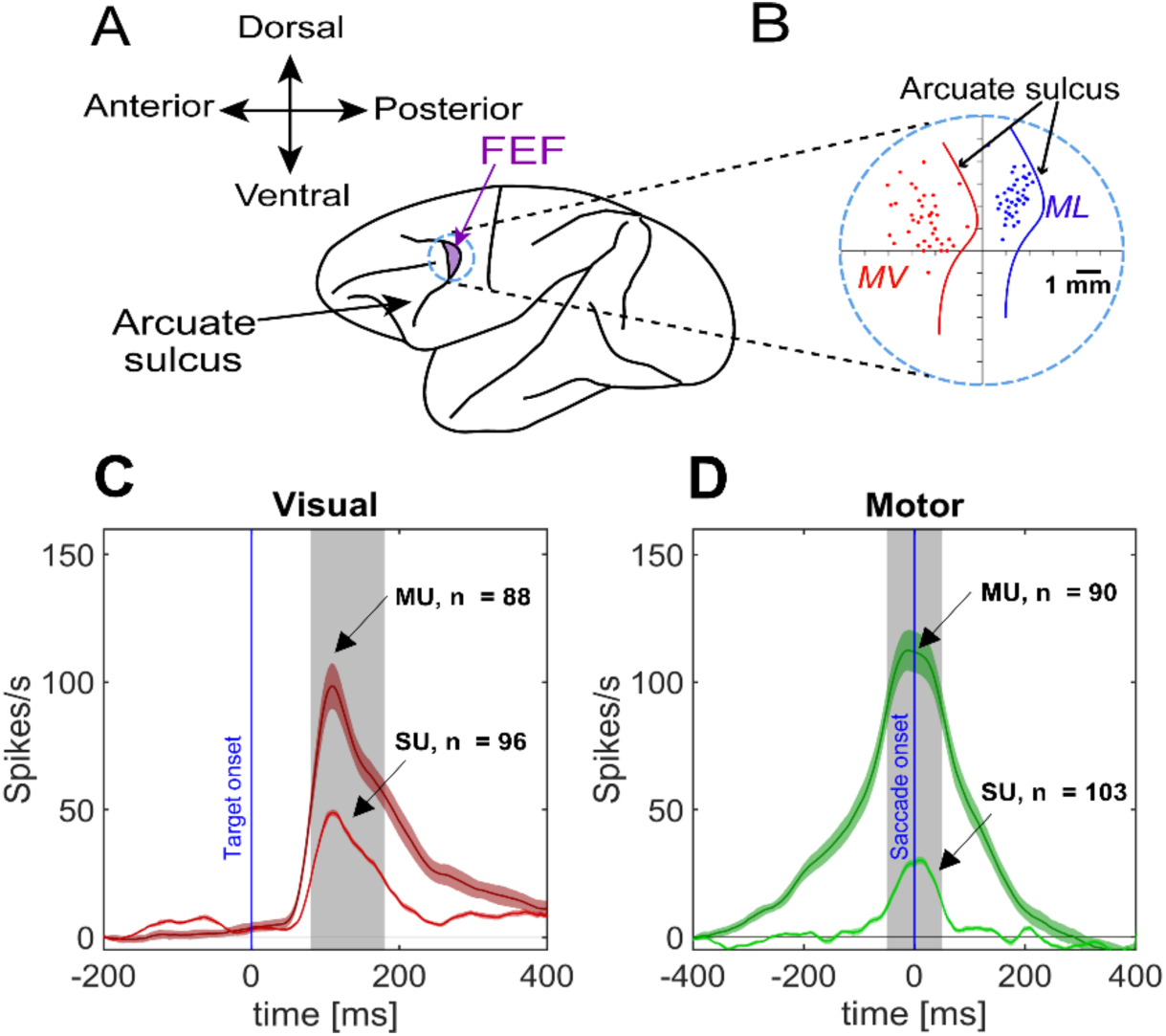
FEF recordings. **A,** Acute FEF recordings were done with a single tungsten electrode. **B,** Overlapped sites of neural recordings from two animals [Monkey V, MV and Monkey L, ML]. Curved lines denote the arcuate sulcus. **C,** Top 10 % (with 95 % confidence interval) spike density plots for SU (red) and MU (dark red) activity for visual neurons. Blue line indicates the target onset. **D,** Same as C but for motor neurons. Blue line stands for saccade onset. Note: the top 10 % activity from each neuron was taken and then pooled across all neurons.

The experiment started once a neuron had reliable spiking activity. The visual or motor response field of a neuron was characterized while the animals performed memory-guided gaze shifts. These gaze shifts started from an initial fixation location to a target (which was randomly flashed one-by-one, generally contralateral to the recording site) within an array (that varied between 4 × 4 to 7 × 7 depending on the size and shape of the response field) of targets (5 –10° apart from each other). The initial fixation locations were jittered within a 7-12° window, which increased variability of 3D gaze, eye, head distributions to initial fixation and the displacements. For our analysis, this variation allowed for separation between effectors and the space-fixed and eye-centered frames of reference for target and gaze (Keith et al., 2009; Sajad et al., 2020). It should be noted that there was no correlation between the initial gaze location and final gaze errors.

The 3D coil signals were sampled at 1000 Hz to measure eye and head orientation in space (Bharmauria et al., 2021, 2020). Gaze (the 2D pointing direction of the eye in space) and eye-in-head positions were derived from these signals (Crawford et al., 1999). When conducting eye movement analysis, the saccade onset (eye movement in space) was marked as the point in time when the gaze velocity increased above 50°/s, while the gaze offset was marked as the point in time when the velocity decreased below 30°/s. The head movement was marked from the saccade onset up to the point in time when the head velocity decreased below 15°/s.

### Single-(SU) and Multi-unit (MU) Activity Sorting

We manually isolated the SU activity from the saved neural recordings using template matching and further confirming it with principal component analysis (PCA). We only carried forward those neurons that had a relatively constant template throughout recording. As the MU cluster comprises different spikes, it generally has a lower amplitude than the individual spikes sorted from it. We re-thresholded the high-pass filtered raw data to isolate the MU activity from the 40 kHz signal with a spike detection threshold of 3.5 standard deviations (SDs). This procedure led us to spiking activity recorded from several neurons recorded within a vicinity of ∼100–200 μm around the electrode tip, thereby allowing us to discriminate the spikes above the noise level (Bullock, 1997; Burns et al., 2010; Buzsáki, 2004; Land et al., 2013; Mattia et al., 2010).

### Data Inclusion Criteria and Sampling Window

Only neurons (or MU sites) that were clearly isolated and task-modulated were analyzed i.e., SU / MU with clear visual activity and / or with perisaccadic movement **(Fig. 2C-D**). Neurons with only post-saccadic activity (activity after the saccade onset) or that lacked significant spatial tuning (see ‘testing for spatial tuning’ below) were excluded from the analysis. While trials where monkeys landed their gaze within the reward acceptance window were included, we excluded trials when the gaze end points went beyond 2° distance from the average gaze endpoint for a given target. In conducting neural activity analysis, the “visual epoch” referred to a fixed 100 ms window of 80-180 ms aligned to the target onset, and the “movement period” referred to a high-frequency perisaccadic 100-ms (−50 - +50 ms relative to saccade onset) window (Sajad et al., 2015). This allowed us to have a good SNR ratio for neuronal activity analysis, and most likely corresponded to the epoch during which gaze shifts influenced neural activity.

### Fitting Neural Response Fields Against Spatial Models

In order to test between different spatial models, they must be experimentally separable (Keith et al., 2009; Sajad et al., 2015). For instance, in our paradigm, the variability induced by memory-guided gaze shift end-points allows one to discriminate target coding from the gaze coding, natural variations in initial eye and head locations allow one to distinguish between different egocentric reference frames, and variable eye and head components for the same gaze shift permit the separation of different effectors (Crawford et al., 1999; Keith et al., 2009). In our method, we use these experimentally derived measures as coordinate systems to analyze neural response field data and then discriminate which spatial model best describes the data.

The logic of our response field fitting in different reference frames is schematized in **Figure 3A**. If the response field data is plotted in the correct / best coordinate system, this should yield a uniform distribution of data with low residuals (i.e., errors between the actual data and a mathematical fit made to these data) **(Fig. 3A**, left). On the other hand, if the coordinate system / fit does not match the data well, this will lead to higher residuals **(Fig. 3A**, right). For example, if an eye-fixed response field is computed in eye-coordinates, this should yield lower residuals compared with head or space coordinates (Sajad et al., 2015). **Figure 3B** represents one head-unrestrained trial (left).

**Figure 3:**
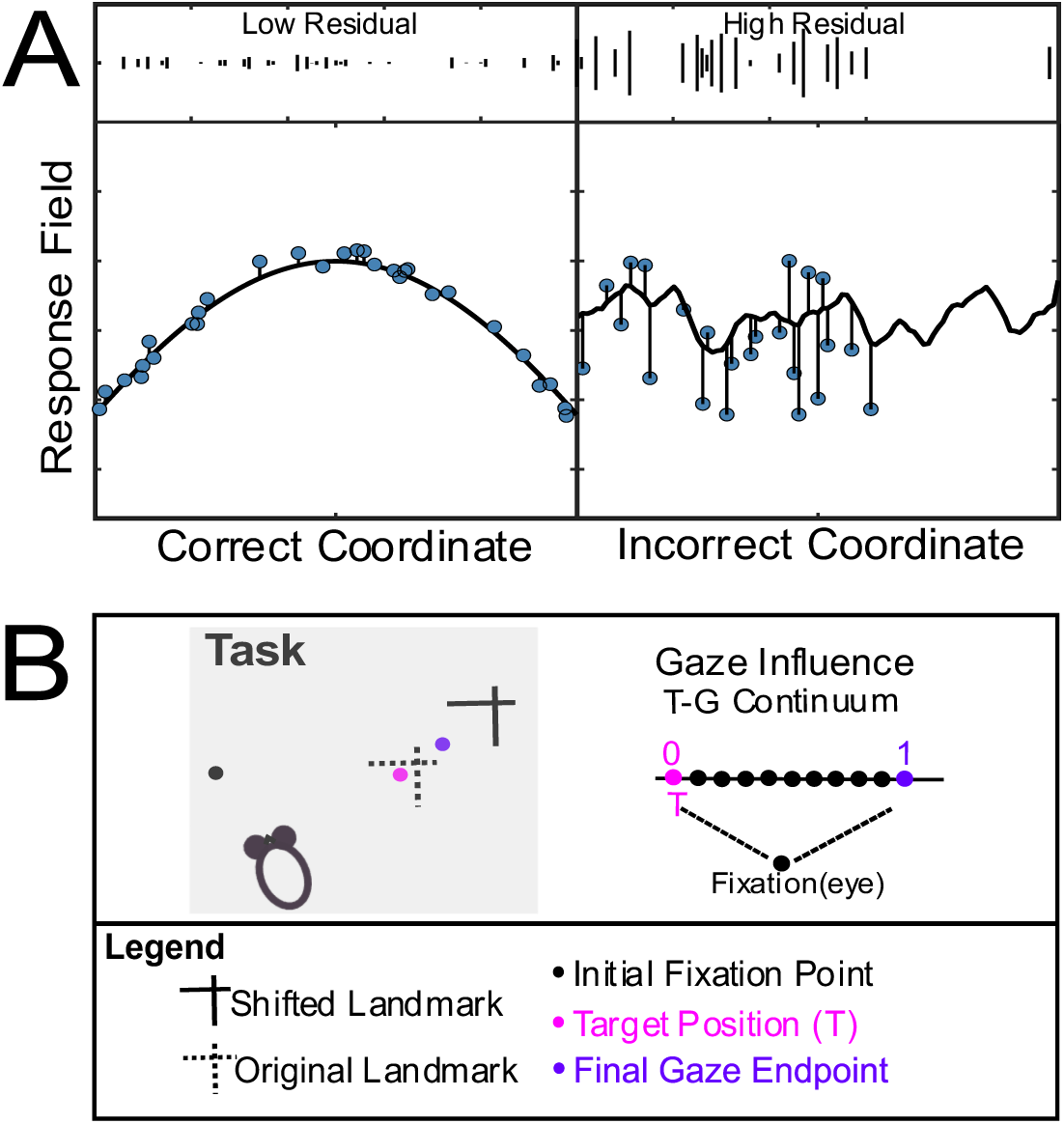
Schematic representation of spatial model fitting procedure. **A,** Illustration of the spatial model fitting technique. The X-axis is the coordinate frame, and the Y-axis is the associated activity to the target. If the activity (blue data points) related to a fixed target is plotted in the correct coordinate frame (left), it yields lower residuals. On the other hand, activity that is plotted in an incorrect coordinate frame, leads to higher residuals (right). **B,** Illustration of the task and gaze influence for a single trial. The black dot depicts the projection of an initial fixation point, and the pink dot represents the target position (T) in relation to the original landmark (L) position (dotted intersecting lines). The solid intersecting lines correspond to the shifted landmark (L’) position and the purple dot depicts the final gaze endpoint (G). An egocentric T-G continuum was plotted to compute the gaze influence by designating the space between T and G into ten uniform steps.

For the actual analysis, we employed a non-parametric fitting method to characterize the neural activity with reference to a specific spatial coordinate system and varied the spatial bandwidth (kernel 2-25°) of the fit to plot any response field size, shape, or contour (Keith et al., 2009). We tested between various spatial models using Predicted Residual Error Some of Squares (PRESS) statistics. In this method, the residual for a trial is computed by comparing the actual activity relative to fits computed from other trials (similar to cross-validation). For a given neuron and temporal window (**Fig. 2**), this was done for all trials, for each bandwidth of fit and for each spatial model tested (e.g., Te, Ge etc.). The model and bandwidth that yields the lowest residuals was deemed ‘best.’ The PRESS residual of this model was then compared with the PRESS residuals of the other models at the same kernel bandwidth using a two-tailed Brown-Forsythe test to test for significant differences in the goodness of fit (typically, a significantly superior fit is only found in a fraction of individual neurons). Finally, we performed the same statistical analysis (Brown-Forsythe) at the population level by comparing the means of PRESS residuals across neurons (DeSouza et al., 2011; Keith et al., 2009). (Significance is more often reached at this level.).

### ‘Cardinal’ Models Tested

We first repeated the model-fitting analysis of the previous study (Bharmauria et al., 2020) on SU and MU activity populations to confirm if Te and Ge were the overall best ‘canonical’ models to represent response field activity in this task. [Note that in the recent investigation (Schütz et al., 2023) these models were called *T_F(e)_* / *G_F(e)_* to specify the Foveal coordinate origin (0,0), but this can be assumed here]. Again, these are not theoretical models, but rather models that were derived from the experimental measures described above. Possible egocentric models included target position *vs.* effector (gaze, eye, head) displacement or position in three egocentric frames (See **supplementary Figure 1** for schematic representations). Specifically, Target in space, eye or head coordinates (Ts, Te, Th); future gaze in space, eye, and coordinates (Ge, Gh, Gs); eye displacement (dE), i.e., the difference between initial and final eye orientation relative to the head; and head displacement (dH), i.e., the difference between initial and final head orientation relative to space; and finally, future orientation of the head in space coordinates (Hs). (We removed dG because it was indistinguishable from Ge in this dataset). We did not test allocentric models (e.g., Target relative to landmark, Landmark relative to Eye) here because they were statistically eliminated at the population level in previous studies (Bharmauria et al., 2021, 2020; Schütz et al., 2023) and our goal here was to test the population egocentric visuomotor transformation.

### Intermediate Spatial Models: the TG continuum

Previous studies suggested that FEF responses do not exactly fit against spatial models like Te or Ge, but actually may fit best against intermediate models between the canonical ones (DeSouza et al., 2011; Sadeh et al., 2015; Sajad et al., 2016, 2015). The spatial continuum between Te and Ge (‘T-G continuum’) provided the best overall description of the data and a signature for the sensorimotor transformation, where G includes variable errors relative to T (Sajad et al., 2015). This continuum was created by taking ten (10 %) steps between and beyond Te and Ge (**Fig. 3B**). Here, these T-G fits were then done at each of these steps, and where the fit yields the lowest PRESS residuals, was considered ‘best’, as described above.

Note that even the best model would not be expected to have zero residuals, because of uncontrolled factors such as biological / measurement noise, non-spatial factors such as arousal / motivation levels (Sajad et al., 2020), and in the case of the motor response, dynamics of the movement (van Opstal, 2023).

### Testing for Spatial Tuning

The method described above works under the assumption that neuronal activity is structured as spatially tuned response fields. Other neurons might implicitly sub-serve the overall population code (Bharmauria et al., 2016a, 2014; Chaplin et al., 2018; Goris et al., 2014; Leavitt et al., 2017; Pruszynski and Zylberberg, 2019; Zylberberg, 2018), but here we were testing for explicit codes. Therefore, we confirmed the spatial tuning of neurons before including them in further analysis. We tested for spatial tuning by randomly (100 times to obtain random 100 response fields) shuffling the firing rate data points across the position data that we obtained from the best model. The mean PRESS residual distribution (PRESS_random_) of the 100 randomly generated response fields was then statistically compared with the mean PRESS residual (PRESS_best-fit_) distribution of the best-fit model (unshuffled, original data). If the best-fit mean PRESS fell outside of the 95 % confidence interval of the distribution of the shuffled mean PRESS, then the neuron’s activity was deemed spatially selective. At the population level, the percentage of neurons showing significant spatial tuning varied through time. Thus, we removed the time steps where the populational mean spatial coherence (goodness of fit) was statistically indiscriminable from the baseline (before target onset). We then defined an index (Coherence Index, CI) for spatial tuning. CI for an SU / MU site, which was calculated as (Sajad et al., 2015):

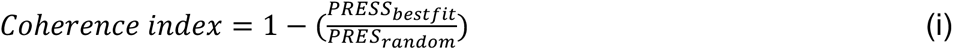

If the PRESS_best-fit_ was similar to PRESS_random_ then the CI would be roughly 0, whereas if the best-fit model is a perfect fit (i.e., PRESS_best-fit_ = 0), then the CI would be 1. We only included those SUs / MU sites in our analysis that showed significant spatial tuning.

### Time-normalization for Spatiotemporal Analysis

In this task, the time between the onset of the target and the gaze onset was not equal across trials because of a variable delay period. To accommodate this variable trial-time for transformations between these events, we time-normalized the duration of each trial (Bharmauria et al., 2021; Sajad et al., 2016). Accordingly, the neural activity across different trials (between the events that we were interested in — 80 ms after the target onset until the onset of saccade for whole population in **Figure 9**) was divided into 14 equal half overlapping bins (with a range of 112.26 – 178.93 ms depending on the trial). If we aligned the trials in a standard way (relative to visual stimulus onset or saccade onset), this would lead to the loss and/or mixing of neural activities across trials. The rationale behind the bin number choice was to make sure that the sampling time window was wide enough, and therefore robust enough, to account for the stochastic nature of spiking activity. During this normalization procedure, the duration of the mask was also convolved in time (on average from step 4.69 to step 7.63). Therefore, the time-normalization procedure was performed to account for the different delay period and the details of this procedure are provided in the previous study (Sajad et al., 2016). For this analysis, the neural firing rate (in spikes/second; the number of spikes divided by the sampling interval for each trial) was sampled at 14 half overlapping time windows from these time-normalized data. The choice for sampling window numbers was based on the approximate ratio of the duration of the visual burst to delay period to movement burst (Sajad et al., 2016). The final (14th) time step in **Figure 9** also contained some part of the perisaccadic sampling window as mentioned above. This time-normalization procedure allowed us to consider the entire time-period (across different trials) of visual–memory– motor responses as an aligned spatiotemporal continuum.

### Code Availability Statement

All codes are custom-written and available on request.

## RESULTS

In the current study, we compared sensorimotor codes by fitting spatial models against SU and MU activity derived from anatomically matched FEF populations. In summary, we recorded task-related neural activity from 257 FEF sites in both animals and isolated a total of 312 FEF units. After applying our exclusion criteria, including tests for spatial tuning, 147 SUs (including visual, V; visuomotor, VM; and motor, M neurons) and the corresponding 101 MU sites, respectively, were taken forward for analysis. Of these, 91 SU sites were accompanied by MU data and 59 MU sites were accompanied by SU data for motor responses (the difference is due to the exclusion criteria). 63 SU sites were accompanied by MU data and 52 MU sites were accompanied by SU data for visual responses. The anatomic distributions of the SU and MU sites were nearly identical in both animals (**Fig. 2B**). Finally, of the 147 SU sites, we analyzed 102 visual and 109 motor responses; and of the 101 MU sites, we analyzed 88 visual and 90 motor responses.

### Summary of Overall Statistical Analysis of Egocentric Models

As a preliminary step, we first tested the Te and Ge models (the overall preferred models for sensory and motor activity in our previous study) against other potential egocentric models (target, eye, head, and gaze displacement or position coding in various egocentric frames of reference). Again, these tests are based on finding the model that yields the overall lowest residuals between our non-parametric fits and actual response field data, and the testing for significance relative to other models.

**Figure 4** summarizes this result for both our SU (**Fig 4 A-B)** and MU (**Fig. 4C-D**) populations (see **Figures 5 and 6** below for example response field fits). The overall patterns for the SU and MU populations were similar: the visual response best codes for Te and the motor response best codes for Ge, as found previously in the SU activity study (Bharmauria et al., 2020). Data points below the horizontal line (P = 0.05) indicate models with worse fits. In the case of the visual responses, all other models were significantly eliminated in both population datasets. In the case of the motor responses, all models were significantly eliminated except for gaze displacement (dG, which is very similar to Ge in this dataset). This does not mean that these populations do not code for other variables in individual cells or in more subtle ways (Schütz et al., 2023) but shows that Te and Ge provide the best overall measures of their explicit population coding scheme. These results confirm the previous SU analysis and tend to suggest that MU populations show similar results. To further investigate this, a more in-depth analysis was performed based on the T-G continuum between the two best models.

**Figure 4:**
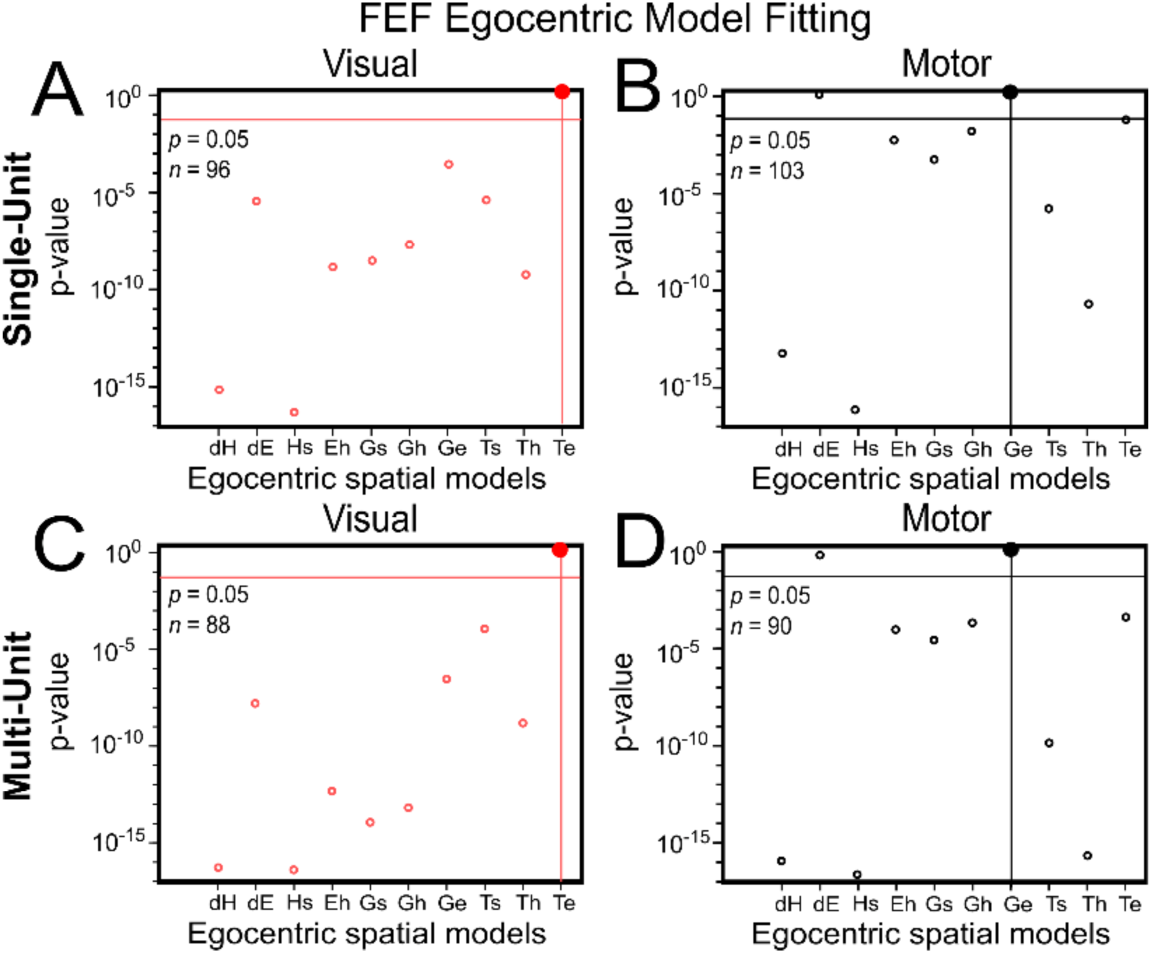
P-Value statistics performed on the residuals of different egocentric models (Brown-Forsythe test) in FEF SU (A) visual and (B) motor responses. Same analysis for MU (C) visual and (D) motor responses. Target in eye (Te) is the best model for the visual response and Ge is the best model for the motor response in both the SU and MU activities. Best model: p = 10^0^ = 1; the best-fit spatial model that yielded the lowest residuals. The horizonal line indicates p = 0.05. Models tested: the difference between the initial and the final head orientation relative to space (dH); the difference between the initial and the final eye orientation relative to the head (dE); Future orientation of the head in space coordinates (Hs); Eye in head (Eh); Future gaze in space (Gs); Future gaze in head (Gh); Future gaze in eye (Ge); Target in space (Ts): Target in head (Th); Target in eye (Te).

**Figure 5:**
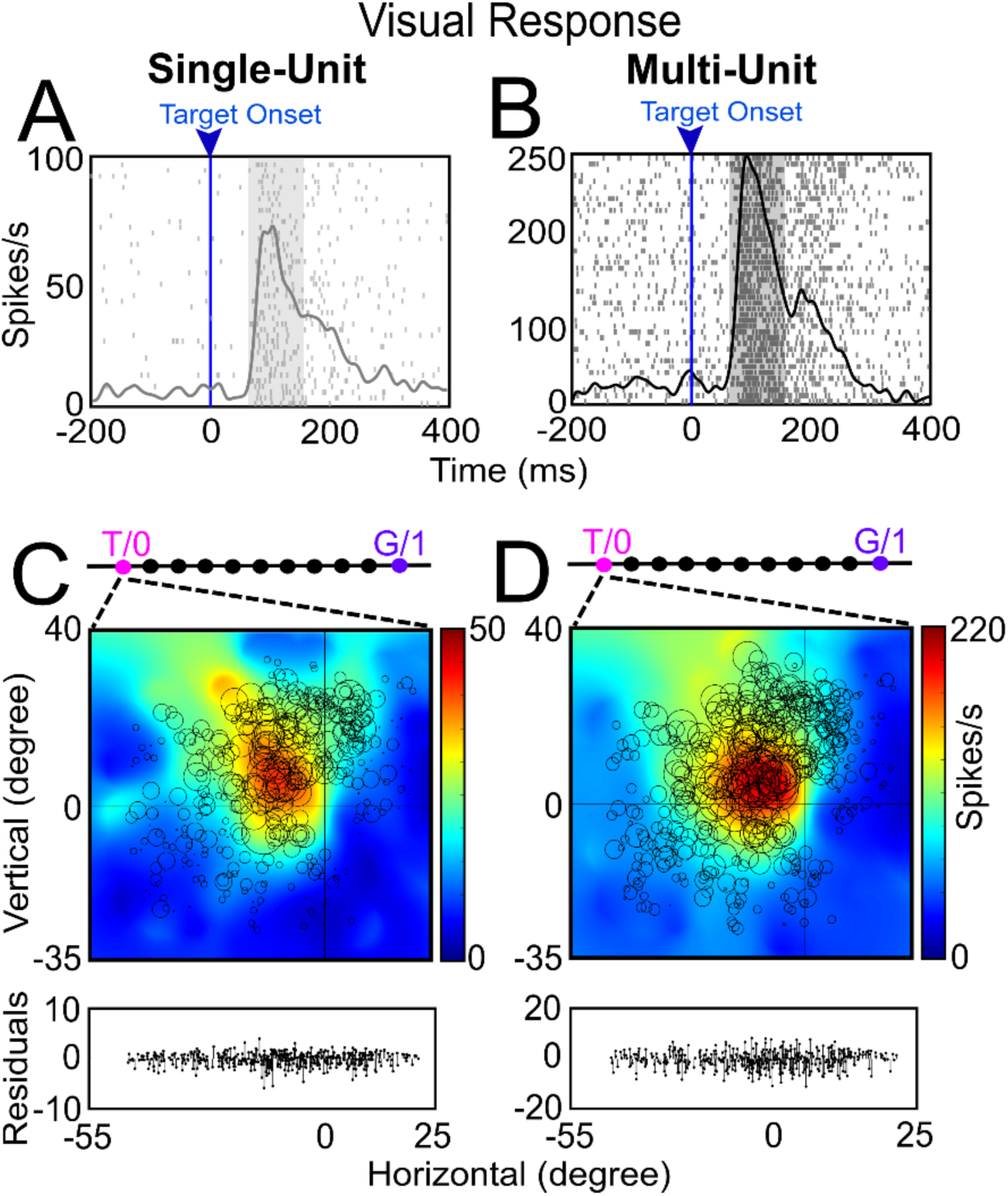
Examples of SU and MU visual response field analysis. **A** and **B**, Raster/spike density plot (with top 10 % of responses) of an SU and MU visual response aligned to target onset (blue arrow), respectively. The shaded gray region represents the sampling window (80-180 ms) for the response field analysis. **C** and **D**, Representation of the response field of SU, and MU visual activity along the T-G continuum, respectively. The size of the circle corresponds to the magnitude of the response and the heat map represents the non-parametric fit to the data (red area is the response field hot spot). The converging dotted lines indicate that both SU and MU fit best at exactly at T. 0, 0 indicates the center of the coordinate system which yielded the lowest residuals (best fit). The graph at the bottom of C and D represents the respective residuals for the response field plots.

**Figure 6:**
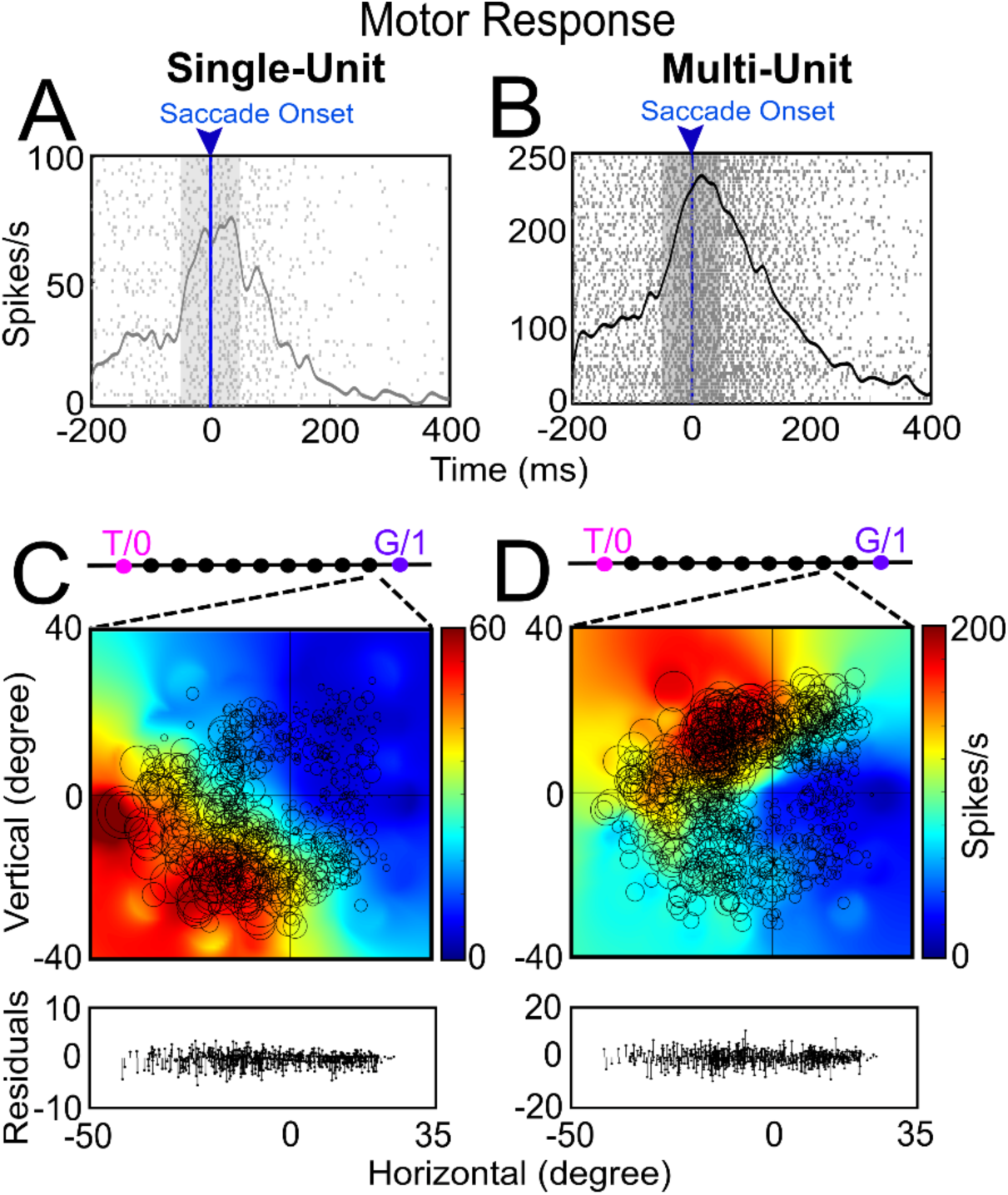
Examples of SU and MU motor response field analysis. **A** and **B**, Raster/spike density plot (with top 10 % of responses) of an SU and the corresponding MU motor response aligned to the saccade onset (blue arrow), respectively. The shaded gray region is the sampling window (– 50-50ms) for the response field analysis. **C** and **D**, Representation of the response field of SU and MU for motor response along the T-G continuum, respectively. The SU response field fits best at the 9^th^ step from T and MU response field fits best at the 8^th^ step. The graph at the bottom of C and D represents the respective residuals for the response field plots.

### Example comparisons of SU and MU activity response fields along the T-G continuum

Previously, it was shown that the major egocentric transformation occurs between target-to-gaze (T-G) by plotting response fields at intermediate steps between a T-G continuum (Bharmauria et al., 2020). Here, a similar analysis on the MU activity was done to compare it with the SU activity. **Figure 5** shows a typical visual response field analysis along the T-G continuum for SU and MU activity recorded at the same site. **Figure 5A** and **B** show the raster and spike density (gray and black curves; representing top 10 % of responses) of an SU and the corresponding MU visual response aligned to the target onset (blue line and arrow), respectively. The shaded gray region is the sampling window (80-180 ms) for the response field analysis. The SU response (firing frequency along the y-axis) is considerably lower than the MU response (because this combined several neurons).

**Figure 5C** shows the best SU fit for the response field along the T-G continuum. The response field best fits at T as indicated by the converging broken lines. Each circle represents the response to one trial, where the bigger the circle, the higher the response is. The heat map represents the non-parametric fit to the data. Note that the main cluster of large circles (and the corresponding red ‘hot-spot’ of the fit) is located in the upper-left quadrant relative to center (0,0, i.e., initial fixation). This localized peak denotes the typical ‘closed’ response field often seen in FEF visual data. The residuals (difference between the fit and each datapoint) are shown below the response fields.

**Figure 5D** shows the response field map of the corresponding MU, again with the best fit at T. In this case, the response fields of SU and MU activity were very similar (both showing similar ‘closed’ organization at the same location), showed the same best fits (T), and their fit residuals were qualitatively similar. Overall, this suggests that SU and MU activity might carry similar spatial information, but we will document this more thoroughly below.

**Figure 6** shows a typical example of an SU and MU motor response field analysis from the same recording site. **Figure 6A-B** shows the raster and spike density (same conventions as **Figure 5** for visual analysis) of an SU (**Fig. 6A**) and MU activity (**Fig. 6B)** for a motor response aligned to the saccade onset (blue line and arrow).

**Figure 6C** shows the response field fit of an SU along the T-G continuum. In this case both the SU and MU response fields showed the typical ‘open’ pattern, with activity continuing to increase away from the center of the coordinates. But in this case, they peaked in different quadrants (upper-left for SU activity, upper-right for MU activity). This suggests that the response field of MU was strongly influenced by other nearby neurons (the second neuron, not shown, isolated from the same site had a response field hot-spot in the second quadrant). However, both the SU and MU activities showed a similar best fit point along the T-G continuum: one step from G (for SU activity) and two steps from G (for MU activity), i.e., close to Ge. In short, despite their differences, both the SU and MU motor responses utilized a similar code (future gaze relative to initial gaze). Collectively, the above examples suggest that MU and SU activities carry comparable information.

### Population Analysis along the T-G continuum

To quantify how representative the above examples were, we plotted the overall distribution of best fits for the SU and MU populations. **Figure 7A1** shows the distribution of best fits for the SU visual responses (n = 96) along the T-G continuum with a primary peak around T and a secondary smaller peak near G. Most units (81.25 %) showed a best fit below 0.5 (i.e., closer to T than G) and conversely,18.75 % showed a best fit closer to G. Overall, the population distribution showed a best fit around T, but was significantly shifted toward G (mean = 0.13; median = 0.10; p = 0.03, one sampled Wilcoxon signed rank test). This suggests that the SU visual responses primarily encoded the target, but some units already predicted the future gaze location.

**Figure 7:**
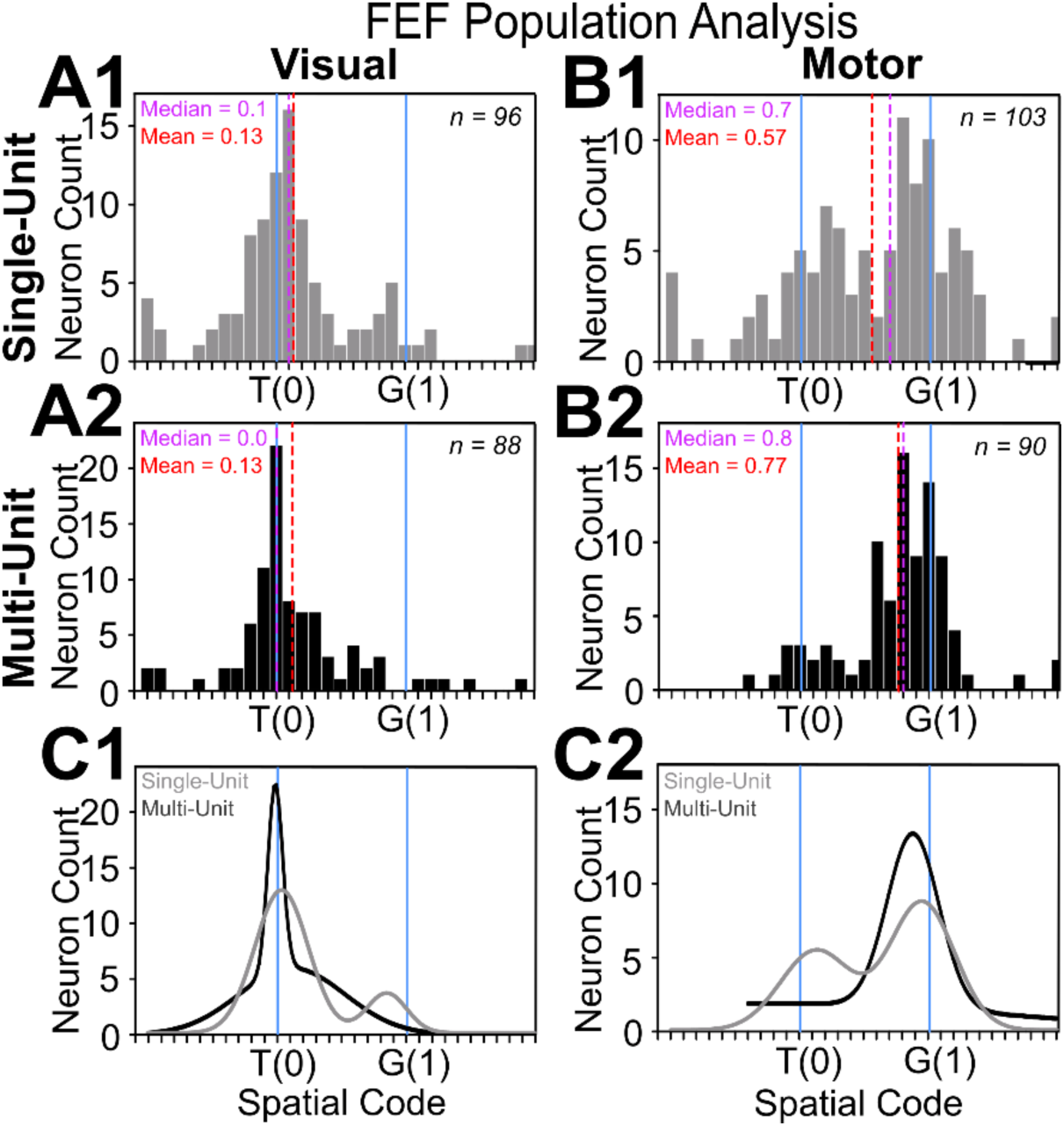
Frequency distribution of SU and MU along the T-G continuum at the population level. **A1**, Frequency distribution of all spatially tuned SU visual responses (n = 96) with the best fit closer to T (mean = 0.13; median = 0.1). **A2**, Frequency distribution of all spatially tuned MU visual responses (n = 88) with the best fit closer to T (mean = 0.13; median = 0). **B1**, Frequency distribution of all spatially tuned SU motor responses (n = 103) with a shifted distribution toward G (mean = 0.57; median = 0.7). **B2**, Frequency distribution of all spatially tuned MU motor responses (n = 90 with a significantly shifted distribution toward G (mean = 0.77; median = 0.8). **C1**, The sum of two Gaussian distributions of SU (two tuning curves) and MU (one tuning curve) visual responses. **C2**, The sum of two Gaussian distributions of SU (two tuning curves) and MU (one tuning curve) motor responses. In both cases, the smaller secondary peak disappeared.

In comparison, the MU visual distribution (n = 88, **Fig. 7A2**) retained the primary peak near T. Although the overall shift from T toward G was still significant (mean = 0.13, median = 0; one sampled Wilcoxon signed rank test p = 0.013), the secondary ‘G’ peak in SU disappeared in the MU fits. Likewise, the number of best fits closer to G than T decreased to 15.9 %. Overall, these results suggests that the MU visual response, comprising several closely connected neurons, better reflected target position whereas gaze prediction was attenuated.

**Figure 7B1-B2** show the similar distributions for motor responses. Overall, the SU motor distribution was significantly shifted toward G compared to the SU visual distribution (p < 0.0001, Mann-Whitney U-test). As expected, the SU motor (n = 103) distribution had a primary peak near G, but still had considerable secondary peak near T. 44.6 % SU motor responses showed best fit scores below 0.5 (closer to T than G), suggesting considerable retention of target information in the motor response. As a result, the overall SU distribution was significantly shifted from both T and G toward an intermediate point (mean = 0.57; median = 0.7; p < 0.0001, one sampled Wilcoxon signed rank test).

The MU motor distributions (n = 88, **Fig. 8B2**) also showed a significant shift toward G (p < 0.0001, one sampled Wilcoxon signed rank test) and a strong ‘G’ peak, but the secondary peak was attenuated: now only 17.77 % of the fits were closer to T than G. As a result, the overall shift in MU (mean = 0.77; median = 0.8) was larger than the SU shift (mean = 0.57; median = 0.7). In other words, MU activity possessed a better canonical gaze code than SU activity but retained less information about the original target location.

**Figure 8:**
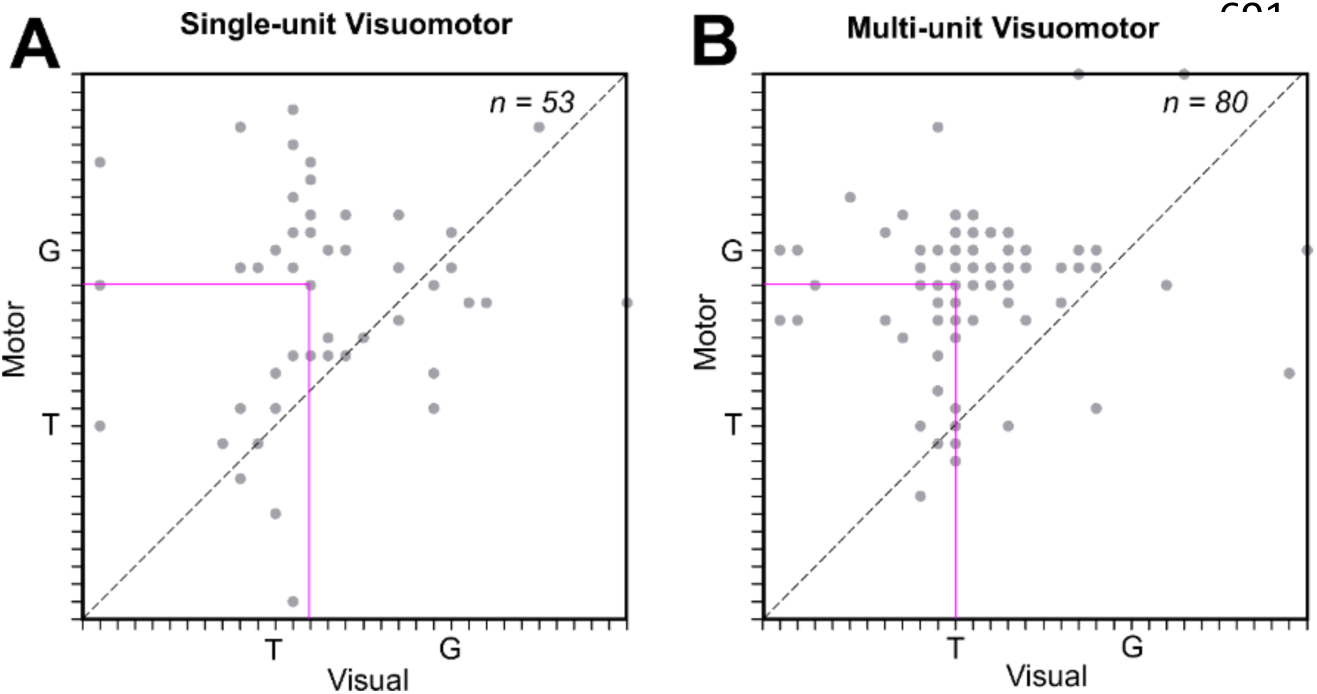
Visuomotor transformation at the single and multi-unit level for visuomotor responses along the T-G continuum. Movement best fit (y-axis) as a function of the corresponding visual best fit (x-axis) for VM neurons (A) and corresponding MU activity (B) sites. Each dot represents one neuron/site. The pink intersecting lines indicate the median of the best-fit for the movement against the median of the best-fit for visual responses. The distribution was significantly shifted toward G in both cases (p < 0.0001, Wilcoxon matched-pairs signed rank test) suggesting visual-to-motor transformation from the target (T) to gaze (G) coding in both cases. However, their (A, B) slopes and elevations were not significantly different from each other, implying a similar profile of visual-to-motor transformations.

**Figure 7C1-C2** superimposes the SU and MU Gaussian distributions for the visual and motor fit frequency distributions shown above. Overall, there was no significant difference between the entire SU and MU distributions for either visual (p = 0.922, Mann-Whitney U test) and motor responses (p = 0.063, Mann-Whitney U test). However, the variances of the fit distributions were significantly different for SU vs. MU motor comparison (F = 2.19, p = 0.0002). Further, in both cases (visual and motor responses), the primary peak (T for visual, G for motor) is similar for SU and MU, but the secondary peak (G for visual, T for motor) is attenuated or missing in the MU population. Consistent with this, the proportion of SU fits below 0.5 (closer to T) in the motor response was significantly less than that of the equivalent MU fits (z-test for proportions; z = 3.63, p = 0.0028). Collectively, these data support the notion that MU visual and motor activities encode purer representations of T and G respectively, whereas the secondary SU codes (predictive G in the visual response and retained T in the motor response) were attenuated.

It is possible that the overall population shift from target to gaze coding could occur only due to transference of activity from visual to motor cells, in SU activity this also occurs within cells as well (Sadeh et al., 2015; Sajad et al., 2015). To test if MU activity shows the same behavior, we contrasted visual and motor codes for neurons/sites that have both visual and the motor responses (**Fig. 8**). **Figure 8A** shows the motor best fits (y-axis) as a function of the visual fits (x-axis) for individual neurons. **Figure 8** illustrates the motor best fits (y-axis) as a function of the visual fits (x-axis) for the MU sites. An upward / leftward shift in these data suggests a transformation from target to gaze coding *within* visuomotor cells (Sadeh et al., 2015; Sajad et al., 2015). In both cases (SU and MU activity), most of the points and the intersection of medians (pink lines) were above the line of unity, and this was significant at the population level (p < 0.0001, Wilcoxon matched-pairs signed rank test) suggesting a visual-to-motor transformation in both cases. Further, the slopes (F = 0.23, p = 0.88) and elevations (F = 0.70, p = 0.40) for SU and MU distributions were not significantly different from each other, suggesting that SU and MU activities reveal similar within-cell / site visuomotor transformation.

### Site-matched Anatomic Comparison: Multi-unit *vs.* Single-unit

We next tested if these fit distributions were comparable for anatomically matched recording sites. Note that the anatomic variance could arise from both 2D distribution of recordings as well as cortical layer depth, which we could not measure accurately. For this analysis only specific sites that contained both SU and MU fits were obtained for visual and / or motor responses, taking the values the SU fits against the MU fits. From **Figure 7** we hypothesized that SU and MU sites might show some anatomical relation, since there was no significant difference in their overall distributions. We did not find significant correlation (**Supplementary Fig. 2**) for MU vs. SU for visual responses for anatomically matched sites (Spearman R = 0.05, Slope = - 0.04 ± 0.08, p = 0.66). However, a modest (Spearman R = 0.20, Slope = 0.10 ± 0.06) but insignificant (p = 0.053) correlation was observed for motor responses. Overall, this suggests that variations in the SU and MU fits do not reflect any anatomical variance in spatial coding.

### Spatiotemporal Analysis

Finally, we performed the 14-step time-normalized analysis (Bharmauria et al., 2020) from visual response until the saccade onset (**Fig. 9**) for both SU and MU activity (see methods). This procedure permitted us to track the progression of T-G along the spatiotemporal domain. The gray curve represents the SU transformation along time (Bharmauria et al., 2020), whereas the black curve corresponds to the visuomotor transformation in the MU activity. In both cases, there was a gradual transition from T-G, however, in the case of the MU activity, the transition seems much ‘smoother’ than the SU, without the dips and rises seen during and after visual mask presentation. Specifically, in the SU activity, the code moves back and forth several times after time-step 6, but in the MU activity, the code is mostly maintained from time-step 8 (post-landmark shift) until the 14^th^ step when the gaze was just imminent. Although subtle differences between the SU and MU data are evident at certain steps between SU and MU, there was no significant difference between the curves (p > 0.05, Mann-Whitney U-test).

**Figure 9:**
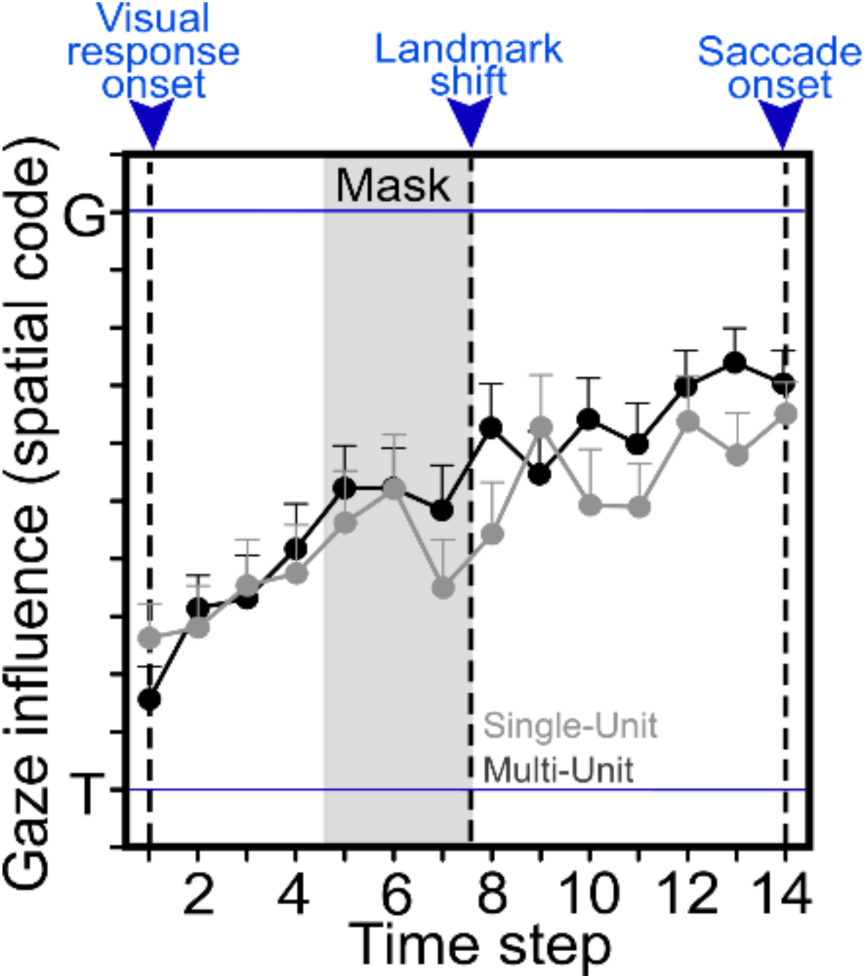
Spatiotemporal analysis at the population level. Progression of the spatial code (gaze influence, mean ± SEM) along the T-G continuum with time (at different time steps) from visual response onset until the saccade onset. There was a gradual and significant progression toward G for both SU (gray) and MU (black) activity.

## DISCUSSION

The aim of this study was to compare sensorimotor transformations derived from single-unit (SU) activity vs. multi-unit (MU) activity from the same FEF recording sessions, using data from a signal-rich memory-delay gaze task. Our analysis of visual and motor response fields (**Figs. 4-6**) results indicated that both SU and MU activity show a fundamental progression from target coding in the visual response to gaze coding in the motor response, except that MU activity does not reveal secondary characteristics, i.e., motor prediction in the sensory response and target location retention in the motor response (**Fig. 7**). This basic transformation was also evident *within* single SU / MU recording sites that showed both visual and motor activity (**Fig. 8**). Finally, SU and MU activity showed a similar temporal target-to-gaze progression during the memory delay, but the transition in MU activity was somewhat ‘smoother’ (**Fig. 9**). One might conclude that MU activity provides an accurate measure of the overall sensorimotor transformation in our dataset (in some ways more clearly than SU activity), whereas SU activity contains additional information that might prove important for more cognitive aspects of the task.

### Multi-unit *vs.* Single-unit Activity

Recently, there has been increased interest in comparing SU and MU activity in both animals and humans (Ayar et al., 2023; Ahmadi et al., 2021; Burns et al., 2010; Christie et al., 2015; Drebitz et al., 2019; Gilja et al., 2015; Pandarinath et al., 2017; Trautmann et al., 2019). For example, Trautmann and colleagues (Trautmann et al., 2019) suggested that MU recordings provide both the practical benefit eliminating spike sorting (required for SU analysis) and reducing the dimensionality of the signal without losing important information content. They performed this analysis on three sets of monkey data from different labs (Ames et al., 2014; Churchland et al., 2012; Kaufman et al., 2014) and reached the same conclusion. This fits with the theory of random projections from high-dimensional statistics (Dasgupta and Gupta, 2003; Ganguli and Sompolinsky, 2012; Indyk and Motwani, 1998; Lahiri et al., 2016), i.e., to recover the geometrical properties of a low-dimensional manifold embedded in a high-dimensional space, one can bypass the coordinates of the high-dimensional space. The results of our study tend to agree with this conclusion, with certain caveats that we will discuss in more detail below.

### Implications for Sensorimotor Control

As noted in the introduction, oculomotor scientists have spent many years studying the transformation from visual target coding to the coding of saccade metrics in saccades and head-unrestrained gaze shifts (Bruce and Goldberg, 1985; Everling and Munoz, 2000; Gandhi and Katnani, 2011; Hanes et al., 1995; Sadeh et al., 2015; Sajad et al., 2015; Schall, 2015; Umeno and Goldberg, 2001). Visual and motor tuning have been dissociated by introducing various position or memory-related errors (Gnadt et al., 1991; Heusser et al., 2022; Mays and Sparks, 1980; Umeno and Goldberg, 2001; White et al., 1994), training animals to make ‘antisaccades’ opposite to the stimulus (Everling and Munoz, 2000) or introducing an intervening saccade (Baizer and Bender, 1989; Camalier et al., 2007). Some of these approaches introduce ‘top-down signals that might alter the circuitry involved (Gaymard et al., 1998; Munoz, 2002; Munoz and Everling, 2004; Pierrot-Deseilligny et al., 2002), but more recent model-fitting and decoding analyses confirm a transition from target-to-saccade coding in SC (Sadeh et al., 2018, 2015), FEF (Bharmauria et al., 2020; Sajad et al., 2016, 2015), and SEF (Bharmauria et al., 2021) activity, even in reactive gaze shifts to visual targets (Sadeh et al., 2020).

The current study shows that both SU and MU activity retain this target-to-gaze transformation, even in the presence of a relatively complex stimulus array. A previous analysis of the same dataset showed that some visual responses also encoded landmark locations (Schütz et al., 2023) and the motor response was modulated by landmark shifts (Bharmauria et al., 2020) but overall, Te to Ge was retained (Bharmauria et al., 2020). Here, we showed that MU activity shows the same sensorimotor transformation (in some ways ’cleaner’, providing a ‘purer’ measure of target and gaze coding at the population levels). Further, whereas the spatiotemporal progression of the SU data showed various dips and rises during the delay period, perhaps related to microsaccades (Heusser et al., 2022), the MU data showed a ‘smoother’ transition, suggesting a more robust sensorimotor signature. Considering the relative ease of recording online MU activity (without the need for offline spike sorting), this suggests that MU activity could be sufficient for certain practical applications requiring low-dimensional information (Ayar et al., 2023; Trautmann et al., 2019). For example, MU activity could be observed to rapidly assess the health of sensorimotor transformations during presurgical recordings in patient populations.

### Implications for the Cognitive Aspects of Movement Control

Higher dimensional neural data from single electrodes may require pooling across several simultaneously recorded MU channels to confidently estimate the neuronal population dynamics (Gao and Ganguli., 2015; Trautmann et al., 2019). Here, the secondary Ge code that was observed in some SU visual responses and the secondary Te code that was observed in some motor responses (Bharmauria et al., 2020) was attenuated for MU analysis. Specifically, these secondary codes are dominated by the primary codes in population statistics (**Fig. 4)** and likewise, they appeared to be ‘drowned out’ by the primary codes of most neurons in the MU activity. Ge activity in the visual response could reflect internal noise that ends up influencing variable errors in final gaze position (Sajad et al., 2020, 2016), but it has been suggested that the prefrontal cortex is involved in the intentional prediction of future gaze errors (Fu et al., 2023). Conversely, the retention of Te in the motor response could be used for motor learning, updating perception and/or corrective saccades (Dash et al., 2015; Lundqvist et al., 2018; Mohsenzadeh et al., 2016; Sajad et al., 2020; Wagner et al., 2021).

**Figure 10** shows a schematic circuit that could explain how these signals arise in SU responses during a memory-delay gaze paradigm, simplifying the process into serial visual, memory, and motor circuits (these signals are both mixed and distributed in the real brain). The sensory input to the visual circuit leads to predominantly eye-centered spatial tuning (red circles) (Sajad et al., 2020; Mullette-Gillman et al., 2005; Snyder, 2005). Some neurons (blue) may already predict future gaze (Ge), while some poorly connected untuned neurons (grey) (Barth and Poulet, 2012) also contributing to population dynamics (Bharmauria et al., 2016a; Levy et al., 2020; Pruszynski and Zylberberg, 2019; Zylberberg, 2018). These gaze-tuned neurons (Ge) in the visual circuit presumably receive feedback input from upstream memory and motor circuits, exhibiting a strong gaze prediction (Masselink and Lappe, 2021; Sajad et al., 2020; Stavisky et al., 2017). But as noted in this study, the latter is strongly attenuated in MU activity.

**Figure 10:**
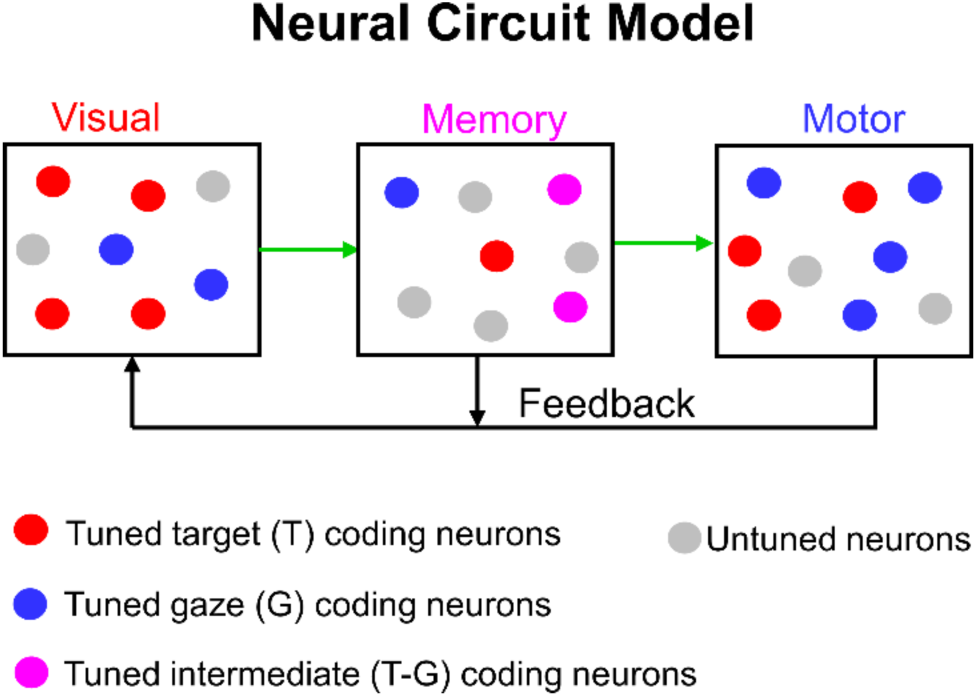
A neural circuit model for SU activity in our task. Visual neurons mostly encode the Target-in-eye coordinates (T; red); however, some neurons (blue) may predict Gaze-in-eye coordinates, possibly due to continuous feedback from memory and motor circuits. Some neurons (grey) remain untuned, although their role in population dynamics cannot be overruled. The visual neurons pass information to memory circuits. Based on previous results (Sajad et al. 2020), memory activity may code target / gaze / intermediate frame (between T and G) coding neuron. This activity is passed on to motor circuits that predominantly code for G, but some neurons retain T. In comparison, MU activity tends to emphasize the main codes at each step and suppress secondary activity.

The visual circuit relays information to the memory circuit, which in turn continuously retains and updates multiplexed signals, with T, G and intermediate coding between T-G (pink neurons) (Bharmauria et al., 2020; Sajad et al., 2020; Sajad et al., 2015). This continuous multiplexed updating mechanism then provides feedback to the visual circuit (Chatham and Badre, 2015; Sajad et al., 2020). This circuit may involve a loop between the LIP, dorsolateral prefrontal cortex (DLPFC), and FEF (Christophel et al., 2017; Lundqvist et al., 2018; Pinotsis et al., 2019). Finally, the system may monitor this feedback loop by up / downregulating feedback gains when feedforward mechanisms fail to provide accurate motor predictions (Franklin et al., 2012; Mathew and Crevecoeur, 2021; Shadmehr and Mussa-Ivaldi, 1994; Wolpert et al., 2011). As noted above, some of these signals are likely obscured in the MU response (as it averages the contribution of all neurons in the circuit), resulting in a ‘smoother’ T-G transition (**Figure 9**).

Lastly, when a gaze shift is cued, activity from the memory circuit is transferred to the motor circuits (blue) in a memory-motor transformation (Bharmauria et al., 2020). This has been shown to involve an additional population shift toward Ge coding (Sajad et al., 2016). Differences of Ge from Te may result from internal noise, endogenous inputs (top-down, updating) or exogenous influences from the internal world (Abedi Khoozani et al., 2022; Bharmauria et al., 2020; Schütz et al., 2023). The blue neurons produce the most accurate Ge command and thus potentiate the gaze shift, whereas the grey neurons remain untuned. Importantly, some Te signals (red neurons) have passed through relatively unaffected, providing the system with an untainted visual memory signal for learning or use in subsequent saccades (Pruszynski and Zylberberg, 2019; Zylberberg, 2018; Barth and Poulet, 2012; Bharmauria et al., 2016a). Overall, the ensemble code shows a progressive transition from Te to Ge across the visual-memory-motor circuitry (Sajad et al., 2020). Once again, the latter codes are attenuated in MU activity, with the result that a similar drawing of the MU circuit would look more like a simple transition from red (target coding) to blue (gaze coding) while missing the various nuances.

### General Implications for Population Coding

Neural data is fundamentally noisy, arising both from endogenous biological noise (Faisal et al., 2008) and from methodological constraints. In the cortex, neurons fire sparsely and exhibit emergent properties in relation to a task/stimulus, but not all activated neurons are necessarily part of the emergent network (Barth and Poulet, 2012; Bharmauria et al., 2016b; Buzsáki, 2010; Molotchnikoff et al., 2019; Singer, 2013). However, if a neural population has homogenous properties, i.e., similar tuning curves, averaging the ensemble activity of several neurons might provide a better estimate of the network’s goal (Quian Quiroga and Panzeri, 2009; Trautmann et al., 2019). Thus, this level of analyses may work best for brain areas with homogeneous populations (Panzeri et al., 2015). This suggests some degree of fundamental homogeneity in the signals we recorded here, despite the relative lack of topography in the recording sites. Moreover, it may also provide information about network level changes after brain damage (Chaudhary et al., 2016) or neuroplasticity (Bharmauria et al., 2022; Kohn, 2007).

Even in the presence of dissimilar tuning curves, or underlying decorrelation, one might still accurately estimate neural manifold geometries using MU channels (Panzeri et al., 2015; Quian Quiroga and Panzeri, 2009). For example, the response field of the motor neuron in **Figure 7C** has a hot-spot in the third quadrant, but the second neuron (not shown) recorded from the same site had a hot-spot in the second quadrant. Despite this, when their MU activity was analyzed, it was possible to create a population hot-spot and determine its motor code (Ge). Consistent with our findings, it has been suggested that only extreme anticorrelations between neural tuning curves disrupt accurate estimation of population manifolds (Trautmann et al., 2019).

### Conclusions

The aim of this study was to test what is retained and what is lost in the methodological transition from SU to MU analysis. An advantage here is that we were able to combine a well-defined sensorimotor system, task, and analytic technique to compare signal coding across both situations. Overall, we find that FEF MU activity carries an excellent representation of the overall sensorimotor transformation for memory-guided gaze shifts but loses some nuances that may be important for the more cognitive aspects of the task. We conclude that MU activity has advantages for applications that must prioritize time over sophistication (potentially including presurgical recordings and some brain-machine interfaces), whereas SU remains relevant for tasks and analyses where more subtle secondary signals and modulations are important.

## Acknowledgement

This project was supported by a Canadian Institutes for Health Research (CIHR) Grant and the Vision: Science to Applications (VISTA) Program, which is supported in part by the Canada first Research Excellence Fund. SS was supported by a Natural Sciences and Engineering (NSERC) Undergraduate Student Award. VB and AS were supported by the NSERC/DFG Brain in Action International Research Training Group (IRTG).VB, XY, and HW were supported by CIHR and VISTA. JDC is supported by the Canada Research Chair Program.

## Supplementary figures

**Supplementary Figure 1:**
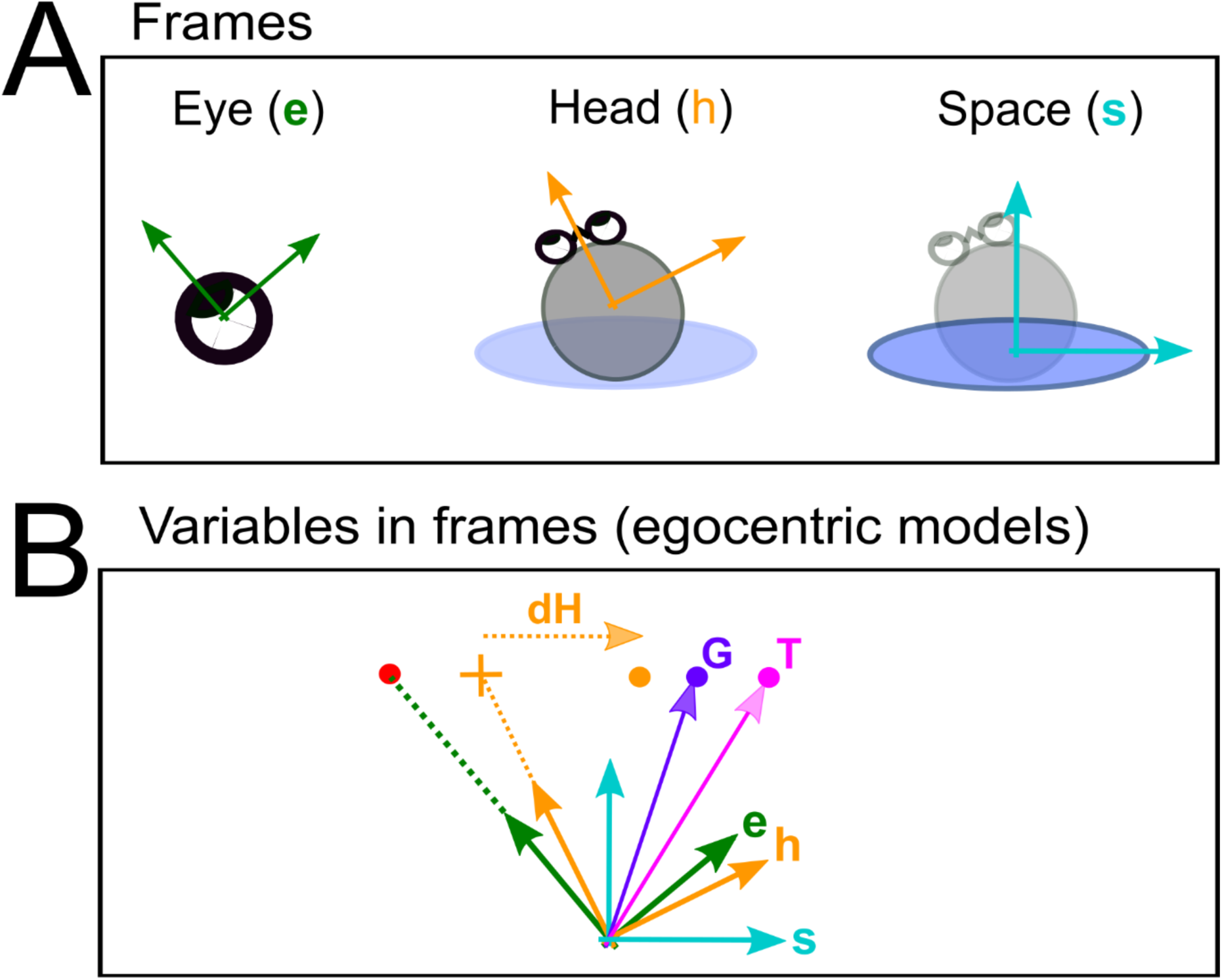
**A**, Basic egocentric reference frames: eye (e), head (h) and body / space (s). **B**, Different variables, and egocentric models for an example trial. Models tested: the difference between the initial and the final head orientation relative to space (dH); the difference between the initial and the final eye orientation relative to the head (dE); future orientation of the head in space coordinates (Hs); Eye in head (Eh); Future gaze in space (Gs); Future gaze in head (Gh) Future gaze in eye (Ge); Target in space (Ts): Target in head (Th); Target in eye (Te).

**Supplementary Figure 2:**
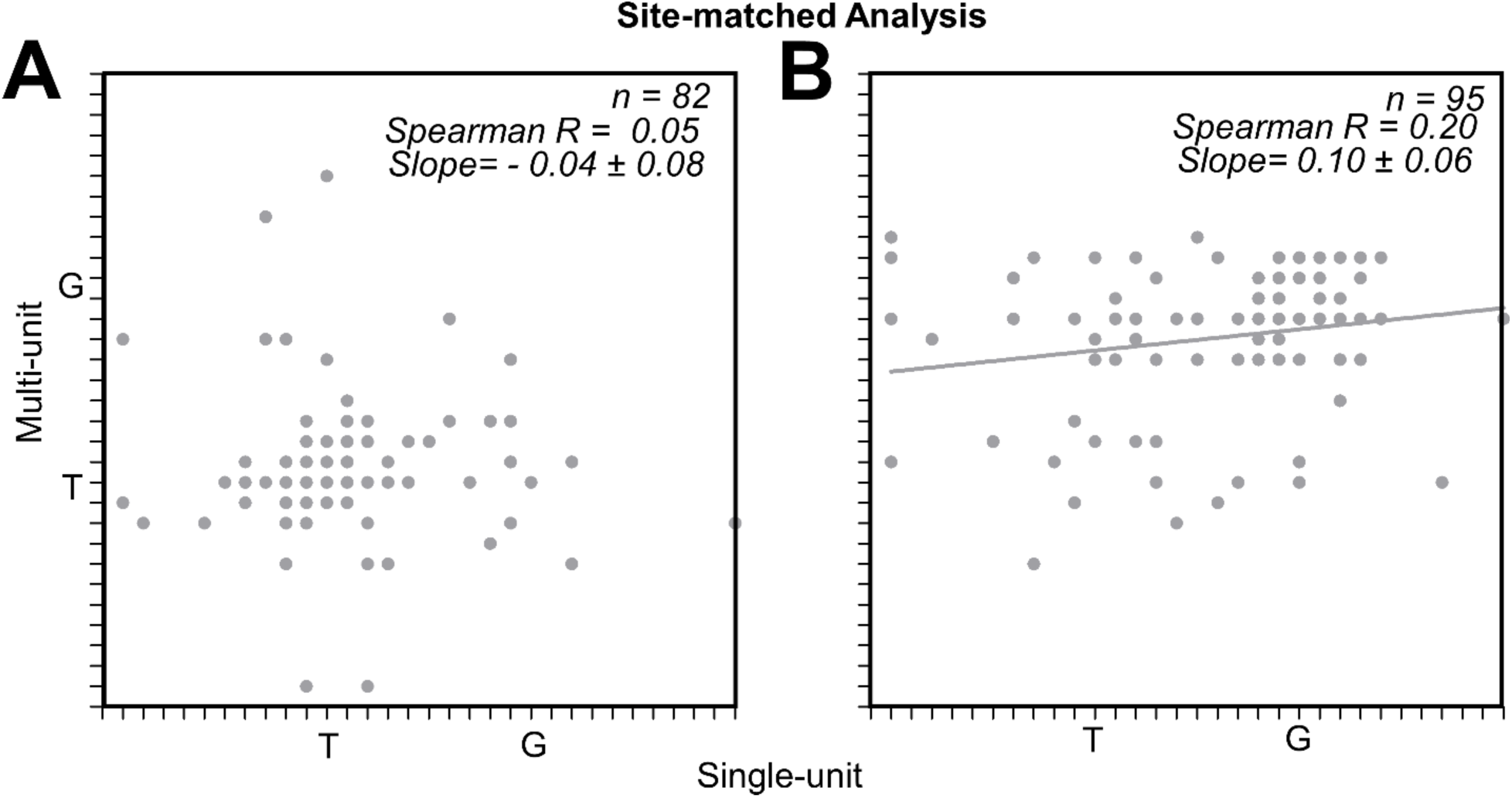
Site-matched analysis. **A**, Visual MU sites as a function of the corresponding best fit score of visual SUs. No significant correlation was observed (Spearman R = 0.05, Slope = - 0.04 ± 0.08, P = 0.66). **B,** Motor MU sites as a function of the corresponding best fit score of motor SUs. A modest correlation (Spearman R = 0.20, Slope = 0.10 ± 0.06) but nearly reaching significance was observed (P = 0.053).

## References

Abedi Khoozani P, Bharmauria V, Schütz A, Wildes RP, Crawford JD (2022) Integration of allocentric and egocentric visual information in a convolutional/multilayer perceptron network model of goal-directed gaze shifts. Cereb Cortex Commun 3:tgac026.

Ahmadi N, Constandinou TG, Bouganis C-S (2021) Inferring entire spiking activity from local field potentials. Sci Rep 11:19045.

Ames KC, Ryu SI, Shenoy KV (2014) Neural dynamics of reaching following incorrect or absent motor preparation. Neuron 81:438–451.

Andersen RA, Brotchie PR, Mazzoni P (1992) Evidence for the lateral intraparietal area as the parietal eye field. Curr Opin Neurobiol 2:840–846.

Andersen RA, Essick GK, Siegel RM (1985) Encoding of spatial location by posterior parietal neurons. Science 230:456–458.

Ayar EC, Heusser MR, Bourrelly C, Gandhi NJ (2023) Distinct context- and content-dependent population codes in superior colliculus during sensation and action. Proceedings of the National Academy of Sciences of the United States of America 120(40):e2303523120

Baizer JS, Bender DB (1989) Comparison of saccadic eye movements in humans and macaques to single-step and double-step target movements. Vision Res 29:485–495.

Barth AL, Poulet JFA (2012) Experimental evidence for sparse firing in the neocortex. Trends in Neurosciences 35:345–355.

Bharmauria V, Bachatene L, Cattan S, Brodeur S, Chanauria N, Rouat J, Molotchnikoff S (2016a) Network-selectivity and stimulus-discrimination in the primary visual cortex: Cell-assembly dynamics. European Journal of Neuroscience 43.

Bharmauria V, Bachatene L, Cattan S, Rouat J, Molotchnikoff S (2014) Synergistic activity between primary visual neurons. Neuroscience 268.

Bharmauria V, Ouelhazi A, Lussiez R, Molotchnikoff S (2022) Adaptation-induced plasticity in the sensory cortex. J Neurophysiol 128:946–962.

Bharmauria V, Bachatene L, Cattan S, Chanauria N, Rouat J, Molotchnikoff S (2016b) High noise correlation between the functionally connected neurons in emergent V1 microcircuits. Exp Brain Research 234:523–532.

Bharmauria V, Sajad A, Li J, Yan X, Wang H, Crawford JD (2020) Integration of Eye-Centered and Landmark-Centered Codes in Frontal Eye Field Gaze Responses. Cereb Cortex 30:4995–5013.

Bharmauria V, Sajad A, Yan X, Wang H, Crawford JD (2021) Spatiotemporal Coding in the Macaque Supplementary Eye Fields: Landmark Influence in the Target-to-Gaze Transformation. eNeuro 8:ENEURO.0446-20.2020.

Brandman DM, Cash SS, Hochberg LR (2017) Review: Human intracortical recording and neural decoding for brain computer interfaces. IEEE transactions on neural systems and rehabilitation engineering : a publication of the IEEE Engineering in Medicine and Biology Society 25:1687.

Bremmer F, Kaminiarz A, Klingenhoefer S, Churan J (2016) Decoding Target Distance and Saccade Amplitude from Population Activity in the Macaque Lateral Intraparietal Area (LIP). Frontiers in Integrative Neuroscience 10:30.

Bruce CJ, Goldberg ME (1985) Primate frontal eye fields. I. Single neurons discharging before saccades. Journal of Neurophysiology 53:603–635.

Bruce CJ, Goldberg ME, Bushnell MC, Stanton GB (1985) Primate frontal eye fields. II. Physiological and anatomical correlates of electrically evoked eye movements. Journal of neurophysiology 54:714–34.

Bullock TH (1997) Signals and signs in the nervous system: the dynamic anatomy of electrical activity is probably information-rich. Proc Natl Acad Sci U S A 94:1–6.

Burns SP, Xing D, Shapley RM (2010) Comparisons of the dynamics of local field potential and multiunit activity signals in macaque visual cortex. J Neurosci 30:13739–13749.

Buzsáki G (2010) Neural syntax: cell assemblies, synapsembles and readers. Neuron 68:362– 385.

Buzsáki G (2004) Large-scale recording of neuronal ensembles. Nat Neurosci 7:446–451.

Camalier CR, Gotler A, Murthy A, Thompson KG, Logan GD, Palmeri TJ, Schall JD (2007) Dynamics of saccade target selection: race model analysis of double step and search step saccade production in human and macaque. Vision Res 47:2187–2211.

Caruso VC, Mohl JT, Glynn C, Lee J, Willett SM, Zaman A, Ebihara AF, Estrada R, Freiwald WA, Tokdar ST, Groh JM (2018a) Single neurons may encode simultaneous stimuli by switching between activity patterns. Nature Communications 9:1–16.

Caruso VC, Pages DS, Sommer MA, Groh JM (2021) Compensating for a shifting world: evolving reference frames of visual and auditory signals across three multimodal brain areas. J Neurophysiol 126:82–94.

Caruso VC, Pages DS, Sommer MA, Groh JM (2018b) Beyond the labeled line: Variation in visual reference frames from intraparietal cortex to frontal eye fields and the superior colliculus. Journal of Neurophysiology 119:1411–1421.

Chaplin TA, Hagan MA, Allitt BJ, Lui LL (2018) Neuronal Correlations in MT and MST Impair Population Decoding of Opposite Directions of Random Dot Motion. eNeuro 5.

Chatham CH, Badre D (2015) Multiple gates on working memory. Current Opinion in Behavioral Sciences 1:23–31.

Chaudhary U, Birbaumer N, Ramos-Murguialday A (2016) Brain-computer interfaces for communication and rehabilitation. Nat Rev Neurol 12:513–525.

Choi Y-S, Koenig MA, Jia X, Thakor NV (2010) Quantifying time-varying multiunit neural activity using entropy based measures. IEEE Trans Biomed Eng 57.

Christie BP, Tat DM, Irwin ZT, Gilja V, Nuyujukian P, Foster JD, Ryu SI, Shenoy KV, Thompson DE, Chestek CA (2015) Comparison of spike sorting and thresholding of voltage waveforms for intracortical brain-machine interface performance. J Neural Eng 12:016009.

Christophel TB, Klink PC, Spitzer B, Roelfsema PR, Haynes J-D (2017) The Distributed Nature of Working Memory. Trends in Cognitive Sciences 21:111–124.

Churchland MM, Cunningham JP, Kaufman MT, Foster JD, Nuyujukian P, Ryu SI, Shenoy KV (2012) Neural population dynamics during reaching. Nature 487:51–56.

Constantin AG, Wang H, Martinez-Trujillo JC, Crawford JD (2007) Frames of reference for gaze saccades evoked during stimulation of lateral intraparietal cortex. Journal of neurophysiology 98:696–709.

Crawford JD, Ceylan MZ, Klier EM, Guitton D (1999) Three-Dimensional Eye-Head Coordination During Gaze Saccades in the Primate. Journal of Neurophysiology 81:1760–1782.

Dasgupta S, Gupta A (2003) An elementary proof of a theorem of Johnson and Lindenstrauss. Random Struct Alg 22:60–65.

Dash S, Yan X, Wang H, Crawford JD (2015) Continuous Updating of Visuospatial Memory in Superior Colliculus during Slow Eye Movements Article Continuous Updating of Visuospatial Memory in Superior Colliculus during Slow Eye Movements. Current Biology 25:267–274.

DeSouza JFX, Keith GP, Yan X, Blohm G, Wang H, Crawford JD (2011) Intrinsic reference frames of superior colliculus visuomotor receptive fields during head-unrestrained gaze shifts. The Journal of neuroscience : the official journal of the Society for Neuroscience 31:18313–26.

Dong Y, Wang S, Huang Q, Berg RW, Li G, He J (2023) Neural Decoding for Intracortical Brain-Computer Interfaces. Cyborg Bionic Syst 4:0044.

Drebitz E, Schledde B, Kreiter AK, Wegener D (2019) Optimizing the Yield of Multi-Unit Activity by Including the Entire Spiking Activity. Frontiers in Neuroscience 13:83.

Edelman GM (1987) Neural Darwinism : the theory of neuronal group selection. Basic Books.

Everling S, Munoz DP (2000) Neuronal correlates for preparatory set associated with pro-saccades and anti-saccades in the primate frontal eye field. Journal of Neuroscience 20:387–400.

Faisal AA, Selen LPJ, Wolpert DM (2008) Noise in the nervous system. Nat Rev Neurosci 9:292–303.

Franklin S, Wolpert DM, Franklin DW (2012) Visuomotor feedback gains upregulate during the learning of novel dynamics. J Neurophysiol 108:467–478.

Fraser GW, Chase SM, Whitford A, Schwartz AB (2009) Control of a brain-computer interface without spike sorting. J Neural Eng 6:055004.

Fu Z, Sajad A, Errington SP, Schall JD, Rutishauser U (2023) Neurophysiological mechanisms of error monitoring in human and non-human primates. Nat Rev Neurosci 24:153–172.

Gandhi NJ, Katnani HA (2011) Motor Functions of the Superior Colliculus. Annu Rev Neurosci 34:205–231.

Ganguli S, Sompolinsky H (2012) Compressed sensing, sparsity, and dimensionality in neuronal information processing and data analysis. Annu Rev Neurosci 35:485–508.

Gaymard B, Ploner CJ, Rivaud S, Vermersch AI, Pierrot-Deseilligny C (1998) Cortical control of saccades. Exp Brain Res 123:159–163.

Gilja V, Pandarinath C, Blabe CH, Nuyujukian P, Simeral JD, Sarma AA, Sorice BL, Perge JA, Jarosiewicz B, Hochberg LR, Shenoy KV, Henderson JM (2015) Clinical translation of a high-performance neural prosthesis. Nat Med 21:1142–1145.

Gnadt JW, Bracewell RM, Andersen RA (1991) Sensorimotor transformation during eye movements to remembered visual targets. Vision research 31:693–715.

Goris RLT, Movshon JA, Simoncelli EP (2014) Partitioning neuronal variability. Nature Neuroscience 17:858–865.

Hafed ZM, Hoffmann K-P, Chen C-Y, Bogadhi AR (2023) Visual Functions of the Primate Superior Colliculus. Annu Rev Vis Sci 9:361–383.

Hanes DP, Thompson KG, Schall JD (1995) Relationship of presaccadic activity in frontal eye field and supplementary eye field to saccade initiation in macaque: Poisson spike train analysis. Experimental Brain Research 103:85–96.

Hebb DO (1949) The Organization of Behavior A NEUROPSYCHOLOGICAL THEORY.

Heusser MR, Bourrelly C, Gandhi NJ (2022) Decoding the Time Course of Spatial Information from Spiking and Local Field Potential Activities in the Superior Colliculus. eNeuro 9:ENEURO.0347-22.2022.

Indyk P, Motwani R (1998) Approximate nearest neighbors: towards removing the curse of dimensionality In: Proceedings of the Thirtieth Annual ACM Symposium on Theory of Computing, STOC ’98, pp604–613. New York, NY, USA: Association for Computing Machinery.

Kao JC, Nuyujukian P, Ryu SI, Shenoy KV (2017) A High-Performance Neural Prosthesis Incorporating Discrete State Selection With Hidden Markov Models. IEEE Trans Biomed Eng 64:935–945.

Kaufman MT, Churchland MM, Ryu SI, Shenoy KV (2014) Cortical activity in the null space: permitting preparation without movement. Nat Neurosci 17:440–448.

Keith GP, DeSouza JFX, Yan X, Wang H, Crawford JD (2009) A method for mapping response fields and determining intrinsic reference frames of single-unit activity: Applied to 3D head-unrestrained gaze shifts. Journal of Neuroscience Methods 180:171–184.

Klier EM, Wang H, Crawford JD (2003) Three-Dimensional Eye-Head Coordination Is Implemented Downstream From the Superior Colliculus. Journal of Neurophysiology 89:2839–2853.

Klier EM, Wang H, Crawford JD (2001) The superior colliculus encodes gaze commands in retinal coordinates. Nature Neuroscience 4:627–632.

Kohn A (2007) Visual adaptation: physiology, mechanisms, and functional benefits. J Neurophysiol 97:3155–3164.

Lahiri S, Gao P, Ganguli S (2016) Random projections of random manifolds.

Land R, Engler G, Kral A, Engel AK (2013) Response properties of local field potentials and multiunit activity in the mouse visual cortex. Neuroscience 254:141–151.

Leavitt ML, Pieper F, Sachs AJ, Martinez-Trujillo JC (2017) Correlated variability modifies working memory fidelity in primate prefrontal neuronal ensembles. Proceedings of the National Academy of Sciences of the United States of America 114:E2494–E2503.

Leuthardt EC, Moran DW, Mullen TR (2021) Defining Surgical Terminology and Risk for Brain Computer Interface Technologies. Front Neurosci 15:599549.

Levy M, Sporns O, MacLean JN (2020) Network Analysis of Murine Cortical Dynamics Implicates Untuned Neurons in Visual Stimulus Coding. Cell Reports 31:107483.

Lewicki MS (1998) A review of methods for spike sorting: the detection and classification of neural action potentials. Network 9:R53–78.

Lewicki MS (1994) Bayesian Modeling and Classification of Neural Signals. Neural Computation 6:1005–1030.

Lundqvist M, Herman P, Miller EK (2018) Working Memory: Delay Activity, Yes! Persistent Activity? Maybe Not. The Journal of neuroscience : the official journal of the Society for Neuroscience 38:7013–7019.

Masselink J, Lappe M (2021) Visuomotor learning from postdictive motor error. eLife 10:e64278.

Mathew J, Crevecoeur F (2021) Adaptive Feedback Control in Human Reaching Adaptation to Force Fields. Frontiers in Human Neuroscience 15.

Mattia M, Ferraina S, Del Giudice P (2010) Dissociated multi-unit activity and local field potentials: a theory inspired analysis of a motor decision task. Neuroimage 52:812–823.

Mays LE, Sparks DL (1980) Dissociation of visual and saccade-related responses in superior colliculus neurons. J Neurophysiol 43:207–232.

Meyers EM (2013) The neural decoding toolbox. Frontiers in Neuroinformatics 7:8.

Miller JK, Ayzenshtat I, Carrillo-Reid L, Yuste R (2014) Visual stimuli recruit intrinsically generated cortical ensembles. Proceedings of the National Academy of Sciences of the United States of America 111:E4053–61.

Mohsenzadeh Y, Dash S, Crawford JD (2016) A State Space Model for Spatial Updating of Remembered Visual Targets during Eye Movements. Frontiers in systems neuroscience 10:39.

Molotchnikoff S, Bharmauria V, Bachatene L, Chanauria N, Maya-Vetencourt JF (2019) The function of connectomes in encoding sensory stimuli. Progress in Neurobiology 101659.

Mtetwa N, Smith LS (2006) Smoothing and thresholding in neuronal spike detection. Neurocomputing, Computational Neuroscience: Trends in Research 2006 69:1366– 1370.

Mullette-Gillman OA, Cohen YE, Groh JM (2005) Eye-centered, head-centered, and complex coding of visual and auditory targets in the intraparietal sulcus. Journal of Neurophysiology 94:2331–2352.

Munoz DP (2002) Commentary: Saccadic eye movements: overview of neural circuitry. Progress in Brain Research 140:89–96.

Munoz DP, Everling S (2004) Look away: the anti-saccade task and the voluntary control of eye movement. Nat Rev Neurosci 5:218–228.

Pandarinath C, Nuyujukian P, Blabe CH, Sorice BL, Saab J, Willett FR, Hochberg LR, Shenoy KV, Henderson JM (2017) High performance communication by people with paralysis using an intracortical brain-computer interface. Elife 6:e18554.

Panzeri S, Macke JH, Gross J, Kayser C (2015) Neural population coding: combining insights from microscopic and mass signals. Trends Cogn Sci 19:162–172.

Park J, Schlag-Rey M, Schlag J (2006) Frames of Reference for Saccadic Command Tested By Saccade Collision in the Supplementary Eye Field. Journal of Neurophysiology 95:159– 170.

Peck CK, Baro JA, Warder SM (1995) Effects of eye position on saccadic eye movements and on the neuronal responses to auditory and visual stimuli in cat superior colliculus. Exp Brain Res 103:227–242.

Pierrot-Deseilligny C, Ploner CJ, Muri RM, Gaymard B, Rivaud-Pechoux S (2002) Effects of cortical lesions on saccadic: eye movements in humans. Ann N Y Acad Sci 956:216– 229.

Pinotsis DA, Buschman TJ, Miller EK (2019) Working Memory Load Modulates Neuronal Coupling. Cerebral Cortex 29:1670–1681.

Pruszynski JA, Zylberberg J (2019) The language of the brain: real-world neural population codes. Current Opinion in Neurobiology 58:30–36.

Quian Quiroga R, Panzeri S (2009) Extracting information from neuronal populations: information theory and decoding approaches. Nature Reviews Neuroscience 10:173– 185.

Quiroga RQ, Nadasdy Z, Ben-Shaul Y (2004) Unsupervised spike detection and sorting with wavelets and superparamagnetic clustering. Neural Comput 16:1661–1687.

Rossi-Pool R, Romo R (2019) Low dimensionality, high robustness in neural population dynamics. Neuron 103:177–179.

Sadeh M, Sajad A, Wang H, Yan X, Crawford JD (2020) Timing determines tuning: A rapid spatial transformation in superior colliculus neurons during reactive gaze shifts. eNeuro 7.

Sadeh M, Sajad A, Wang H, Yan X, Crawford JD (2018) The Influence of a Memory Delay on Spatial Coding in the Superior Colliculus: Is Visual Always Visual and Motor Always Motor? Frontiers in Neural Circuits 12:74.

Sadeh M, Sajad A, Wang H, Yan X, Crawford JD (2015) Spatial transformations between superior colliculus visual and motor response fields during head-unrestrained gaze shifts. European Journal of Neuroscience 42:2934–2951.

Sajad A, Sadeh M, Crawford JD (2020) Spatiotemporal transformations for gaze control, Physiological reports. NLM (Medline).

Sajad A, Sadeh M, Keith GP, Yan X, Wang H, Crawford JD (2015) Visual-Motor Transformations Within Frontal Eye Fields During Head-Unrestrained Gaze Shifts in the Monkey. Cerebral cortex (New York, NY : 1991) 25:3932–52.

Sajad A, Sadeh M, Yan X, Wang H, Crawford JD (2016) Transition from Target to Gaze Coding in Primate Frontal Eye Field during Memory Delay and Memory-Motor Transformation. eNeuro 3.

Schall JD (2015) Visuomotor Functions in the Frontal Lobe. Annual Review of Vision Science 1:469–498.

Schlag J, Schlag-Rey M (1987) Evidence for a supplementary eye field. Journal of Neurophysiology 57:179–200.

Schütz A, Bharmauria V, Yan X, Wang H, Bremmer F, Crawford JD (2023) Integration of landmark and saccade target signals in macaque frontal cortex visual responses. Commun Biol 6:938.

Shadmehr R, Mussa-Ivaldi FA (1994) Adaptive representation of dynamics during learning of a motor task. J Neurosci 14:3208–3224.

Sharma G, Annetta N, Friedenberg D, Blanco T, Vasconcelos D, Shaikhouni A, Rezai AR, Bouton C (2015) Time Stability and Coherence Analysis of Multiunit, Single-Unit and Local Field Potential Neuronal Signals in Chronically Implanted Brain Electrodes. Bioelectron Med 2:63–71.

Siegel M, Buschman TJ, Miller EK (2015) Cortical Information Flow During Flexible Sensorimotor Decisions. Science 348:1352–1355.

Singer W (2013) Cortical dynamics revisited. Trends in Cognitive Sciences 17:616–626.

Smith E, Kellis S, House P, Greger B (2013) Decoding stimulus identity from multi-unit activity and local field potentials along the ventral auditory stream in the awake primate: implications for cortical neural prostheses. J Neural Eng 10:016010.

Snyder LH (2005) Frame-up. Focus on “eye-centered, head-centered, and complex coding of visual and auditory targets in the intraparietal sulcus.” J Neurophysiol 94:2259–2260.

Sommer MA, Wurtz RH (2001) Frontal eye field sends delay activity related to movement, memory, and vision to the superior colliculus. J Neurophysiol 85:1673–1685.

Stavisky SD, Kao JC, Ryu SI, Shenoy KV (2017) Motor cortical visuomotor feedback activity is initially isolated from downstream targets in output-null neural state space dimensions. Neuron 95:195–208.e9.

Telenczuk B, Destexhe A (2020) Local Field Potential, Relationship to Unit Activity In: Encyclopedia of Computational Neuroscience (Jaeger D, Jung R eds), pp1–6. New York, NY: Springer.

Todorova S, Sadtler P, Batista A, Chase S, Ventura V (2014) To sort or not to sort: the impact of spike-sorting on neural decoding performance. J Neural Eng 11:056005.

Trautmann EM, Stavisky SD, Lahiri S, Ames KC, Kaufman MT, O’Shea DJ, Vyas S, Sun X, Ryu SI, Ganguli S, Shenoy KV (2019) Accurate Estimation of Neural Population Dynamics without Spike Sorting. Neuron 103:292–308.e4.

Umeno MM, Goldberg ME (2001) Spatial processing in the monkey frontal eye field. II. Memory responses. J Neurophysiol 86:2344–2352.

van Opstal AJ (2023) Neural encoding of instantaneous kinematics of eye-head gaze shifts in monkey superior Colliculus. Commun Biol 6:927.

Wagner I, Wolf C, Schütz AC (2021) Motor learning by selection in visual working memory. Sci Rep 11:9331.

White JM, Sparks DL, Stanford TR (1994) Saccades to remembered target locations: an analysis of systematic and variable errors. Vision research 34:79–92.

Wolpert DM, Diedrichsen J, Flanagan JR (2011) Principles of sensorimotor learning. Nat Rev Neurosci 12:739–751.

Xing D, Yeh C-I, Shapley RM (2009) Spatial spread of the local field potential and its laminar variation in visual cortex. J Neurosci 29:11540–11549.

Zhang Z, Constandinou TG (2021) Adaptive spike detection and hardware optimization towards autonomous, high-channel-count BMIs. J Neurosci Methods 354:109103.

Zylberberg J (2018) The role of untuned neurons in sensory information coding. bioRxiv 134379.

